# Discovery and Engineering of Retrons for Precise Genome Editing

**DOI:** 10.1101/2024.07.21.604473

**Authors:** Jesse D. Buffington, Hung-Che Kuo, Kuang Hu, You-Chiun Chang, Kamyab Javanmardi, Brittney Voigt, Yi-Ru Li, Mary E. Little, Sravan K. Devanathan, Blerta Xhemalçe, Ryan S Gray, Ilya J. Finkelstein

## Abstract

Retrons are promising gene editing tools because they can produce multi-copy single-stranded DNA in cells via self-primed reverse transcription. However, their potential for inserting genetic cargos in eukaryotes remains largely unexplored. Here we report the discovery and engineering of highly efficient retron-based gene editors for mammalian cells and vertebrates. Through bioinformatic analysis of metagenomic data and functional screening, we identified novel retron reverse transcriptases (RTs) that are highly active in mammalian cells. Rational design further improved the editing efficiency to levels comparable with conventional single-stranded oligodeoxynucleotide donors but from a genetically encoded cassette. Small molecule inhibitors of non-homologous end joining factors and Cas9-DNA repair protein fusions further increase homology-directed repair. Retron editors also exhibited robust activity with Cas12a nuclease and Cas9 nickase, expanding the genomic target scope and bypassing the need for a DNA double-stranded break. Using a rationally engineered retron editor, we incorporate a split GFP epitope tag for live cell imaging. Finally, we develop an all-RNA delivery strategy to enable DNA-free gene editing in cells and vertebrate embryos. This work establishes retron editors as a versatile and efficient tool for precise gene editing, offering new opportunities for biotechnology and biomedical research.

## Introduction

Precise genome editing is a cornerstone of biomedicine and gene therapy. However, installing specific edits via templated homology-directed repair (HDR) is limited by the challenge of delivering and integrating an exogenous template DNA into the genome [1–3]. The most efficient delivery methods are viral vectors and synthetic DNA donors [4–7]. Viral vectors can induce genotoxicity via insertional mutagenesis, are depleted during the gene editing window, and are not suitable for multiplexed applications [5, 7, 8]. Synthetic DNA templates must be transfected or electroporated into cells, do not target the nucleus, and are incompatible with RNA-based delivery in cells and organisms [4, 6]. Therefore, recent gene editing technologies have begun to harness reverse transcriptases (RTs) that can continuously generate the template DNA near its edit site. [9–35]. Among these approaches, retron-RTs are especially promising tools because they are self-priming and can generate high copies of long single-stranded DNAs (ssDNAs) *in vivo*.

Retrons are bacterial anti-phage defense systems that consist of a self-priming reverse transcriptase (retron-RT), a cognate non-coding (ncRNA) that primes & templates reverse transcription, and an accessory protein that participates in anti-viral immunity [36–44]. The ncRNA consists of two main regions: the msr (msDNA-specific region) and the msd (multi-copy ssDNA-coding region) (**Fig. 1A**). The msr is located at the 5’ end of the ncRNA and forms a specific structure that is recognized by the RT [45, 46]. This region typically contains one to three stable stem-loops with 7-10 base pair stems and 3-10 nucleotide loops. The msr also includes a highly conserved guanosine residue at the 5’ end, which serves as the branching point for initiating reverse transcription (**Fig. 1A**). The msd is positioned downstream of the msr and can be divided into two parts: a dispensable region that can be replaced with a desired sequence (i.e., the donor DNA for genome editing) and a conserved region that is essential for the proper folding and function of the ncRNA. The RT primes from the msr and uses the msd as a template [45, 46]. The resulting msDNA remains covalently linked to the ncRNA through a 2’,5’-phosphodiester bond formed between the branching guanosine residue in the msr and the 5’ end of the msDNA [45, 46]. The host RNAse H degrades the RNA-DNA hybrid to expose the ssDNA (**Fig. 1A, right**) [39]. By replacing the dispensable msd region with a user-programmable sequence, retron-RTs can generate templates for homology-directed DNA repair in cells.

**Figure 1:**
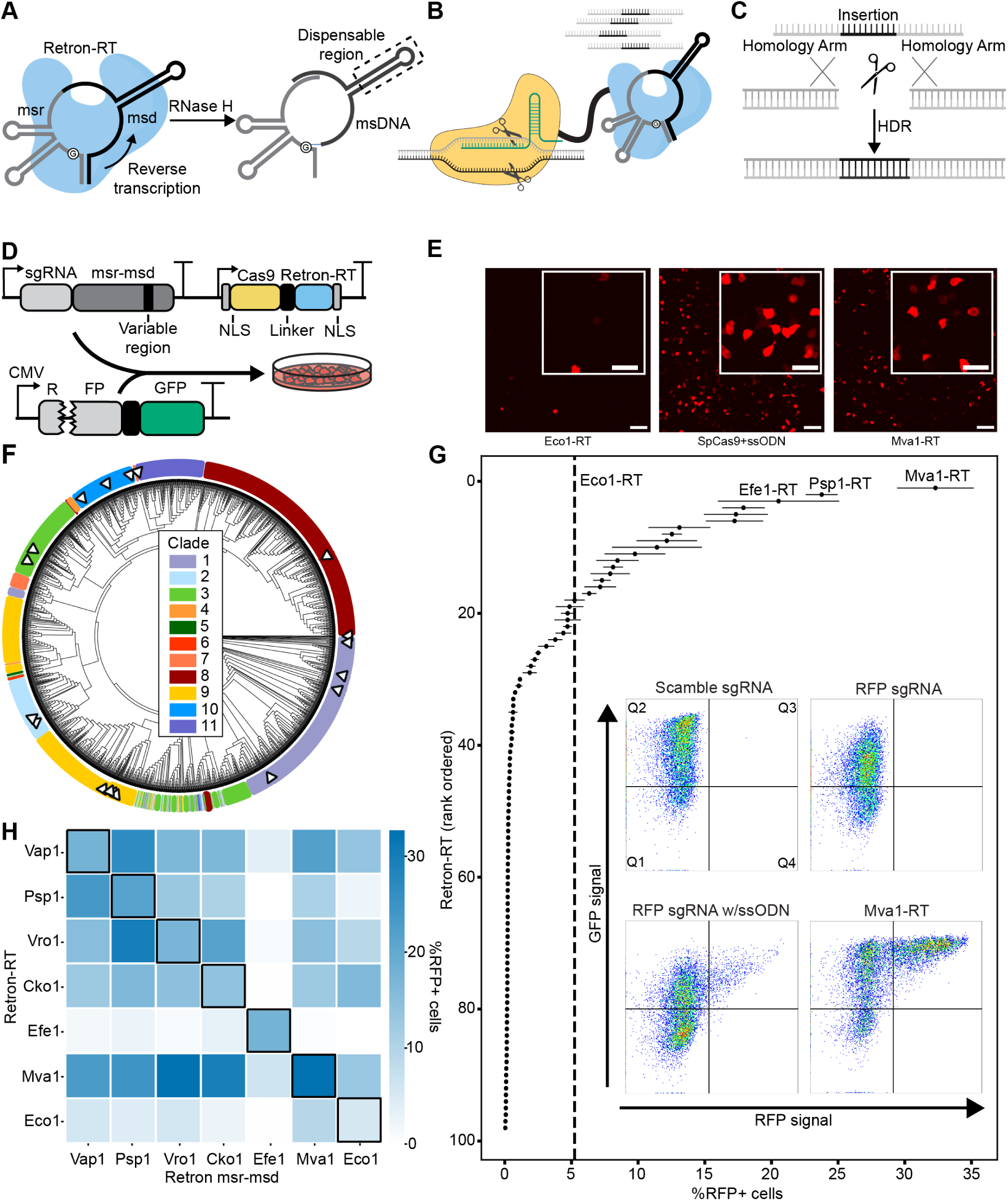
A metagenomic survey reveals highly active RTs in mammalian cells. A) Retron-RTs self-prime from a non-coding RNA, termed the *msr-msd*. msr: gray; msd stem: black; variable region: black, in dotted box. The arrow indicates the direction of reverse transcription. B) Schematic of a retron editor. The RT is linked to Cas9 (shown) or another nuclease. C) Reverse transcription of the variable region of the *msd* generates a ssDNA template for homology-directed repair of the cleavage site. D) A plasmid-encoded fluorescent reporter assay. The RFP has a 9bp deletion proximal to a Y64L mutation to completely turn off RFP fluorescence. The reporter is co-transfected with a plasmid that encodes the retron editor, along with an *msd* that repairs the broken RFP. RFP+ cells are imaged via confocal microscopy and quantified via flow cytometry. E) Confocal microscopy images of cells transfected with Cas9-Eco1-RT (left), Cas9 + ssODN (middle), and Cas9-Mva1-RT (right). F) Phylogenetic classification of novel retron systems discovered from metagenomic sources. These systems are classified into clades, as described in[51]. G) Rank ordered list of RFP repair efficiency with 98 metagenomically discovered retron-RTs using flow cytometry. Dashed line: RFP+ repair with Eco1-RT. Inset: flow cytometry data for Cas9 with a scrambled sgRNA (top left); RFP-targeting sgRNA (top right); RFP sgRNA and a ssODN repair template (bottom left); RFP sgRNA and Mva1-RT (bottom right). Error bars: mean of three biological replicates. The three most active RTs are labeled. H) Gene editing activity of the six most active retron-RTs, along with Eco1-RT with a cognate (diagonal) or non-cognate msr-msd. Flow cytometry was used to score activity with the transient RFP reporter. Heat map: mean of three biological replicates.

Recent studies have demonstrated the potential of retrons coupled with CRISPR-Cas9 to enhance precise genome editing in bacteria, yeast, plants, and mammalian cells [17, 19, 24, 30–32, 34] (**Fig. 1B**). Early studies fused the Cas9 single guide RNA (sgRNA) to a retron ncRNA with a modified donor msd [32]. This increases the local concentration of the donor template at a double-stranded DNA break (DSB), biasing repair toward templated HDR (**Fig. 1C**). Alternative designs fused the retron RT to Cas9 directly with the same ultimate goal [30, 34]. Despite these advances, the potential of retrons for precise genome editing has yet to be fully realized. To date, only a handful of retron RTs have been benchmarked in mammalian cells. Further engineering of the retron editor system, along with the ncRNA, and delivery methods can further improve editing efficiency. Developing a flexible framework for retron editor optimization will greatly expand the utility of this tool across all domains of life.

Here, we report the discovery and engineering of a highly efficient retron gene editor. We bioinformatically identified >500 high-confidence retrons from metagenomic sources. Using a functional reporter system, we screened 98 variants in mammalian cells and identified 17 RTs that were more active than the previously established Eco1-RT. Further rational design achieved editing efficiencies that are comparable to conventional single-stranded oligodeoxynucleotide (ssODN) donors. Steering DNA repair outcomes towards HDR via small molecule inhibitors and Cas9-DNA repair protein fusions boosted targeted DNA in-sertion. Retron editors also function with Cas12a, significantly broadening their genomic target range. The nickase Cas9(D10A) also supports retron editing, and this activity can be improved with DNA repair protein fusions. We apply retron editors for installing in-frame epitopes for live cell imaging in U2OS cells. Finally, we demonstrate all RNA-based retron editing in cell lines and vertebrates. Broadly, this work establishes retron editors as a powerful tool for templated cargo insertion, opening new gene editing modalities.

## Results

### A metagenomic survey identifies retron-RTs active in mammalian cells

We hypothesized that a metagenomic survey of retron-RTs would uncover variants that improve homology-directed repair in heterologous hosts. Towards this goal, we developed a pipeline to phenotypically assess retron-RTs using fluorescent proteins in HEK293T cells (**Fig. 1D**). The reporter expresses RFP and GFP that are separated by a ribosomal skipping T2A sequence [47]. RFP has a 9 base pair (bp) deletion (Δ9) adjacent to a Y64L mutation [48]. These mutations ensure that RFP(Δ9) is dark until the wild type (WT) sequence is restored via templated HDR following a Cas9–generated double-stranded break (DSB). GFP serves as a transfection control and also reports on Cas9-generated insertions and/or deletions (indels) that shift the open reading frame out of frame. As expected, transfecting Cas9 and a single-stranded oligodeoxynucleotide (ssODN) donor restored RFP fluorescence (**Fig. 1E, middle**). The well-characterized Eco1-RT also repaired RFP, although HDR activity was substantially lower than the ssODN (**Fig. 1E, left**). The dispensable msd region included 29 nt of homology flanking a 9 nt insertion that paired with the target strand and also reverted the Y64L mutation. Having established this assay, we proceeded to test additional RTs from diverse microbes.

Retron-RTs are ubiquitous throughout bacteria, but only a handful have been tested experimentally [33]. Therefore, we developed a bioinformatics pipeline to identify new retron-RTs from metagenomic sources (**Fig. S1A**). We first annotated all RTs in the NCBI database of non-redundant bacterial and archaeal genomes, as well as 2M partially assembled bacterial genomes from the human microbiome [49, 50]. We reasoned that human microbiome-derived RTs will also be active under physiological conditions. After identifying likely retron-RTs, we searched for the msr-msd non-coding RNA (ncRNA) and accessory proteins. This search identified >500 high-confidence, non-redundant retrons with well-annotated msr-msds (**Fig. S1B**). We classified these systems into a phylogenetic tree and sorted them into 11 clades following a prior bioinformatic survey (**Fig. 1F**) [51]. The highest-confidence systems across multiple clades were prioritized for experimental characterization in mammalian cells.

Next, we screened 98 retron-RTs using flow cytometry and the RFP(Δ9) reporter (**Fig. 1G**). As expected, transfecting the Cas9-sgRNA plasmid with a ssODN partially restored the RFP signal (**Fig. 1G, inset**). Thirty-one RTs (32% of all tested systems) restored RFP fluorescence (defined as >1% RFP+ cells). Eco1-RT restored RFP+ signal in 5% of the cells. Ten RTs had >2-fold higher repair activity than Eco1-RT (**Fig. 1G**). Mva1-RT, derived from *Myxococcus vastator*, had an editing efficiency that was 6-fold higher than Eco1-RT in this transient repair assay (**Fig. 1G, inset**). The best-performing RTs all belonged to clade 9, indicating that these enzymes are especially active in mammalian cells, and/or that our bioinformatics workflow was most accurate in predicting the msr-msd sequences from this clade. The top systems also showed mostly RFP+ cells via confocal microscopy (**Figs. 1E, S2A**). To test these RTs in a genomic reporter, we also integrated the RFP cassette into the AAVS1 locus of HEK293T cells. While Eco1-RT was minimally able to restore RFP signal, the top six RTs were all active in the genomically integrated assay (**Fig. S2B**). *Escherichia fergusonii* (Efe1)-RT was the most active in the genomic reporter, restoring RFP+ signal *∼*10-fold better than Eco1-RT.

Retron-RTs co-evolve with a cognate msr-msd, but their ability to reverse transcribe from the msr-msd of other retrons is unknown. Therefore, we explored the feasibility of using two or more orthogonal RTs for multiplexed retron editing (**Fig. 1H**). We tested the RFP repair activity of six active novel RTs, along with Eco1-RT, when co-expressed with the msr-msd from other systems. Mva1-RT, the most broadly cross-reactive RT, only shared 33-36% amino acid sequence identity and 49-59% msr-msd nucleotide identity with all other systems (**Fig. S3A**). In contrast, Efe1-RT shared 44-46% amino acid sequence identity but remained exclusive to its cognate msr-msd. The remaining RTs could use a broad range of msr-msds, including those from Eco1-RT. *Vibrio rotiferianus* (Vro1)- and *Vibrio aphrogenes* (Vap1)-RTs showed comparable activity with *Proteus sp.* (Psp1) msr-msd and their native msr-msds. Notably, all RTs shared a similar msr-msd secondary structure, including a palindromic repeat and an extended msd hairpin (**Fig. S3B**). We conclude that retron-RTs are more flexible in msr-msd utilization than previously appreciated and that caution should be taken when combining multiple systems in a single cell line. Further, Efe1-RT is the most active enzyme in the genomic reporter assay and strikes an excellent balance between high activity and specificity for gene editing applications.

Next, we tested the top-performing RTs for their ability to integrate a 10 nt cargo into native genomic loci (**Fig. 2**). Two rounds of PCR were used to generate libraries for next-generation DNA sequencing (NGS) (**Fig. 2A**). The first PCR reaction was primed outside the homology arms to avoid amplifying the reverse transcribed ssDNA. The second PCR reaction barcoded the amplicons for short-read NGS. Cas9 and a ssODN were used as a positive control, and to benchmark the RT fidelity. We distinguished retron editing efficiency from Cas9-generated indels by scoring the percentage of modified reads that had the intended insert relative to all modified reads. Editing efficiencies ranged from 8-30%. Efe1-RT showed the highest editing activity at EMX1 and CFTR, consistent with the genomically integrated RFP(Δ9) assay (**Fig. 2B**). Thus we selected Efe1-RT for all subsequent experiments.

**Figure 2:**
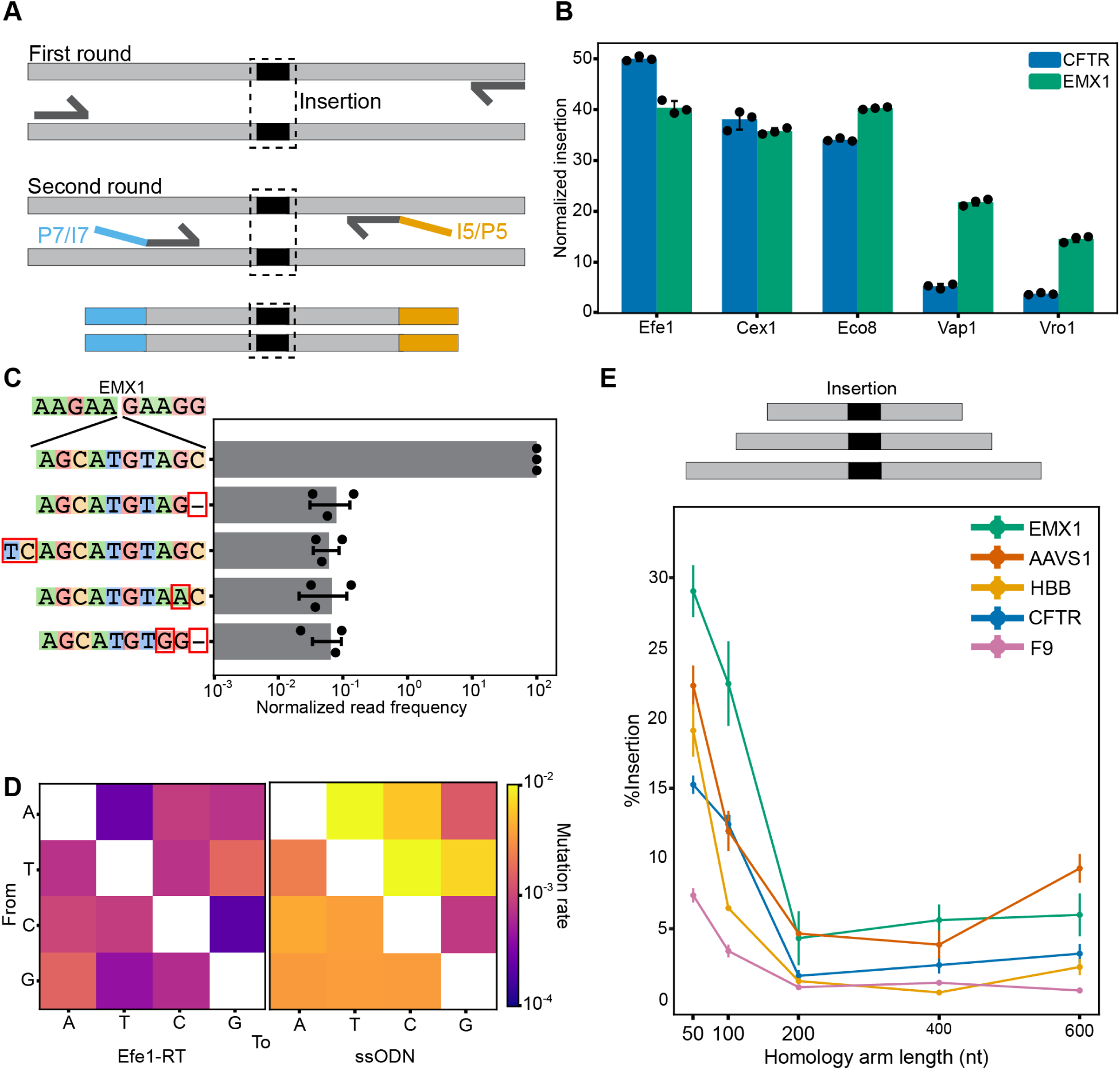
Efe1-RT catalyzes precise genomic insertions across multiple loci. A) Schematic of the NGS library preparation strategy. Genomic DNA is first amplified with primers that are outside the homology arms to avoid amplifying the retron-synthesized msDNA. After gel extraction, a second round of PCR amplifies and barcodes the insertion site for deep sequencing. Blue, orange: universal Illumina P5/I5 and P7/I7 adapters and indices. B) Normalized insertion efficiency for the top 5 retron-RTs at the CFTR and EMX1 loci. Error bars: mean of three biological replicates (dots). C) The relative insertion frequency of a 10 nt cargo at the EMX1 locus, along with the four most frequent misincorporated sequences. The most common errors are a deletion or insertion at the periphery of the homology arms. Error bars: mean of three biological replicates (dots). D) Efe1-RT substitution errors (left) are less frequent than ssODN insertion (right) at the EMX1. Substitution rates are computed from the insert in the EMX1 locus. Heat map: three biological replicates. E) Schematic (top) and results (bottom) of the insertion efficiency with Efe1-RT as a function of the homology arm length. Templated insertion is most active with 50 nt homology arms at five genomic loci. Error bars: mean of three biological replicates (dots).

Deep sequencing of the insert at the EMX1 locus showed that >99% of Efe1-RT driven insertion events contained the intended 10 nt cargo (**Fig. 2C, D**). The most frequent imperfect insertions included indels and single nucleotide substitutions that are the signature of alternative end joining pathways [52, 53]. To determine whether RT fidelity is contributing to this error, we conducted a similar analysis for the Cas9+ssODN experiment at EMX1 (**Fig. 2D**). Efe1-RT error rates were *∼* 10*^−^*^3^ *−* 10*^−^*^4^ (errors/nt), consistent with the substitution rates measured for high fidelity RTs (**Fig. 2D**) *in vitro* [54, 55]. In contrast, ssODN substitution rates were 10-fold greater at the same locus. These results indicate that RT-based editing fidelity exceeds that of conventional ssODNs and that repair fidelity may be limited by cell-intrinsic repair pathways (also see below).

We next tested retron editing across five native loci and as a function of the homology arm length. In all cases, inserting 10 nt was most efficient with 50 nt homology arms (7-28% insertion rates across five loci) (**Fig. 2E**). Editing efficiency was strongly correlated with Cas9 cleavage activity at each locus. Insertion efficiency generally matched, and in the case of EMX1 and HBB, exceeded the edit rates with Cas9 + ssODN. We conclude that short homology arms support the highest insertion rates in our system.

### Rational Engineering of Retron Editors Increases Insertion Activity

To further improve insertion activity, we optimized ncRNA expression, nuclear localization signals (NLSs), and the nuclease-RT linker in an Efe1-RT-based retron editor (**Fig. 3**). The integrated RFP(Δ9) reporter was used for rapid iterative screens. Splitting the sgRNA and msr-msd into two transcripts increased editing efficiency (**Fig. 3A,B**). In the ”split” design, sgRNA expression was driven by a U6 promoter, and the msr-msd was transcribed via the H1 promoter (**Fig. 3A**). These results suggest that fusing the sgRNA and the msr-msd may impact Cas9 and/or RT activity, possibly by misfolding the structural elements of each ncRNA, or by imposing additional steric constraints. All subsequent experiments used the split design.

**Figure 3:**
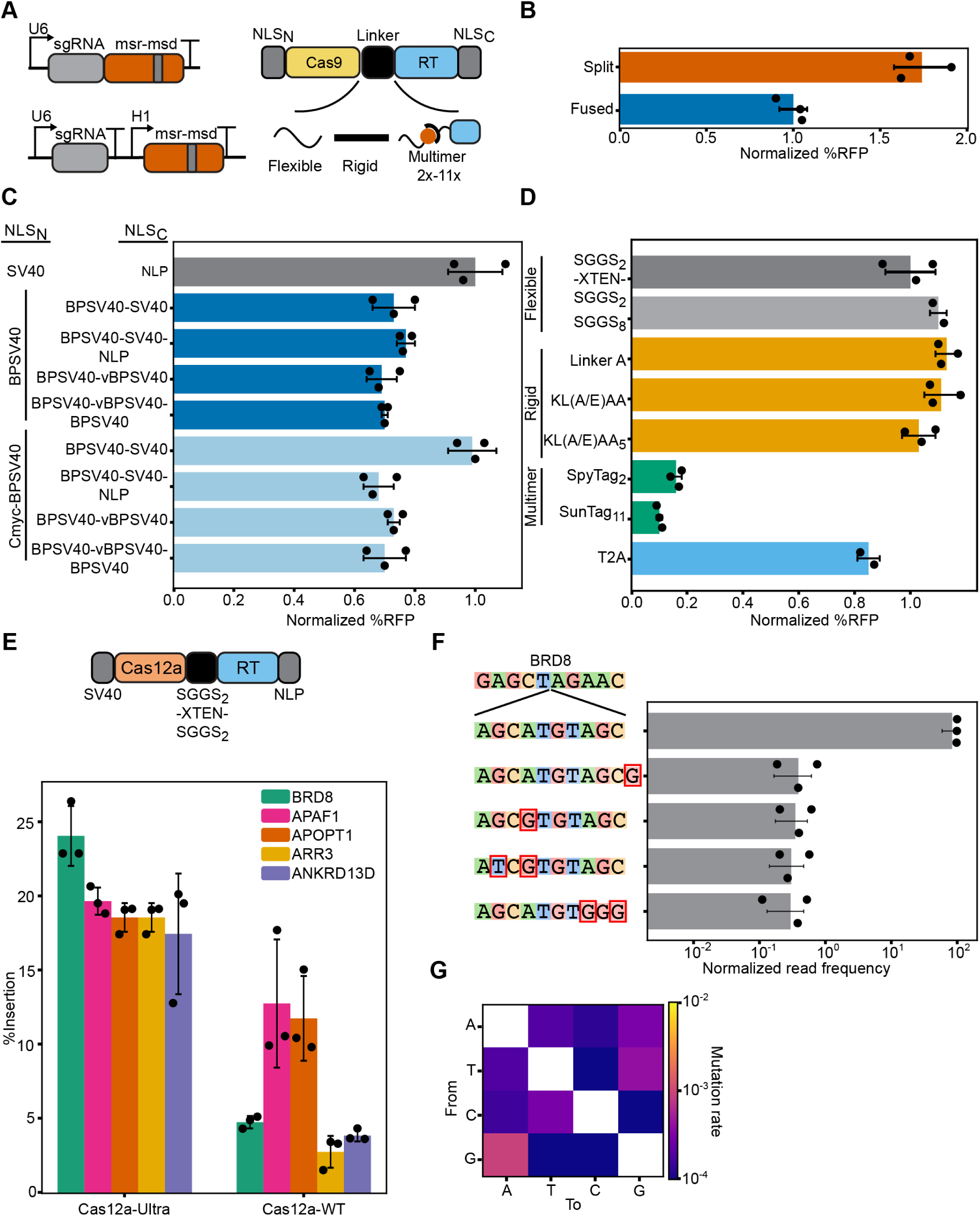
Rational Engineering of an Efe1-RT-based retron editor. A) We tested the effect of splitting the sgRNA and msr-msd (left), the identity of the nuclear localization sequences (NLSs), and the linker between the Cas9 and Efe1-RT (bottom). B) Expressing the sgRNA and msr-msd increased gene editing by 60% relative to a fused sgRNA-msr-msd design. C) Optimization of the N- and C-terminal nuclear localization sequences (NLSs). Gray: reference design that was used for normalization. Error bars: mean of three biological replicates (dots). D) Optimization of the linker peptide between the Cas9 and Efe1-RT. Retron editors tolerate a broad range of flexible (light gray) and rigid (orange) linkers. Splitting the two enzymes via a ribosomal skipping peptide (T2A, light blue) also retains most activity. However, multimerization domains abrogated activity (green). Dark gray: reference design that was used for normalization. Error bars: mean of two or three biological replicates (dots). E) Schematic (top) and editing activity of a Cas12a-based retron editor at five genomic loci. Error bars: mean of three biological replicates (dots). F) The relative insertion frequency of a 10 nt cargo at the BRD8 locus, along with the four most frequent misincorporated sequences. In contrast to Cas9-based retron editors, Cas12a retron editors generate substitution errors in the insert. Error bars: mean of three biological replicates (dots). G) Insert substitution frequencies across five loci. Heat map: mean of three biological replicates.

Nuclear import can be a rate-limiting factor in mammalian gene editing [56–61]. Therefore, we tested 25 N- and C-terminal NLS combinations that were previously reported to improve Cas9-based gene editing (**Fig. 3C & Table 5**). A combination of N- and C-terminal bipartite SV40 (BP-SV40) NLSs showed the highest RFP repair activity. Adding cMyc-SV40 to the N-terminus and two or more SV40s to the C-terminus increased Cas9 cleavage 1.6-fold relative to an N-terminal SV40 and C-terminal NLP signal, as measured by a reduction in GFP+ cells. However, this did not increase templated DNA insertion (**Fig. 3C**). Retron-RTs interact extensively with their ncRNA and msDNA via their C-terminal domains [41, 62]. An extended C-terminal NLS may impair this interaction, reducing overall repair activity, but not Cas9-catalyzed DNA cleavage. We conclude that nuclear import is likely not the limiting factor for templated insertion.

The linker between the nuclease and the RT can also impact editing outcomes. We tested 15 linkers with various physical properties, including a ribosomal skipping T2A sequence between Cas9 and Efe1-RT (**Fig. 3D and Table 6**). Separating the two enzymes via a T2A sequence reduced RFP+ cells by 20% relative to the reference linker, (SGGS)_2_-XTEN-(SGGS)_2_ [63]. Next, we tested a panel of flexible (e.g., (GGS)_N_) and rigid (e.g., (KL(A/E)AA)_n_) linkers. Retron editors accommodated a broad range of linker designs (**Table 6**). However, multimerizing the RT on Cas9 via SpyTags or SunTags reduced RFP+ signal by 85-90% [64]. As Cas9-Sun/SpyTag-Cas9 fusions are reportedly active in mammalian cells, we speculate that multimerization disrupts RT activity [64, 65]. We conclude that retron editors can accommodate a broad range of linker geometries, and even split enzyme designs.

To expand the retron editor target range, we tested Efe1-RT with WT AsCas12a and AsCas12a Ultra at five genomic loci (**Fig. 3E-G**) [66]. Cas12a and Efe1-RT were fused via an (SGGS)_2_-XTEN-(SGGS)_2_ linker, and the crRNA and msr-msd were expressed from independent promoters. In all cases, AsCas12a Ultra-RT fusions had higher insertion activity than WT AsCas12a (**Fig. 3E**). This higher activity is due to the increased cleavage by AsCas12a Ultra relative to WT enzyme (**Fig. S4**). Deep sequencing of the BRD8 locus confirmed that 99% of the inserts had the insert sequence (**Fig. 3F**). The most frequent error was an indel outside of the immediate insert site, followed by mismatches within the insert. Base substitution rates across five genomic loci closely matched the pattern that was observed at EMX1 and were lower than those with Cas9 + ssODN donor (**Figs. 3G, 2D**). In sum, retron editors can be assembled from a broad range of ncRNA, NLS, & linker configurations, and can be paired with Cas9 and Cas12a nucleases to expand their target range.

### Channeling repair pathway choice increases insertion efficiency

DNA repair via non-homologous end joining (NHEJ) limits templated DNA insertion in mammalian cells [53, 67]. To further increase retron editing efficiency, we sought to channel repair away from NHEJ via two approaches (**Fig. 4A**). First, we focused on small molecule inhibitors that inhibit NHEJ or enhance HDR. AZD7648 and M3814 inhibit the DNA-dependent protein-kinase catalytic subunit (DNA-PKcs) to improve templated repair of Cas9 breaks [68–71]. TAK-931 is a CDC7-selective inhibitor that arrests cells in S phase, thereby increasing the HDR time window [72]. We first established the optimal working concentrations for each inhibitor (**Fig. S5A**). All three inhibitors improved insertion activity, with the strongest improvements with AZD7648 at all loci (**Fig. 4B, left**). In contrast, M3814 showed strong improvements at all loci except HBB. Next, we tested whether retron editors can insert larger cargos with 50 nt homology arms, and how this is modulated by inhibitors or DNA repair proteins (**Fig. 4C**). AZD7648 stimulated insertion of 25 and 50 nt inserts by 8.8 and 6.0-fold respectively at EMX1 (**Fig. 4D**). TAK-931 showed more modest 2.4 and 1.8 fold editing increases for 25 and 50 nt cargos respectively. A comparison of Cas9-generated indels and retron-driven insertions confirmed that AZD7648 doesn’t increase Cas9 cleavage, but increases the utilization of a template ssDNA for genomic repair. Further, AZD7648 reduced the mutational signature at the target site, further highlighting the utility of repair pathway modulation in retron editing applications.

**Figure 4:**
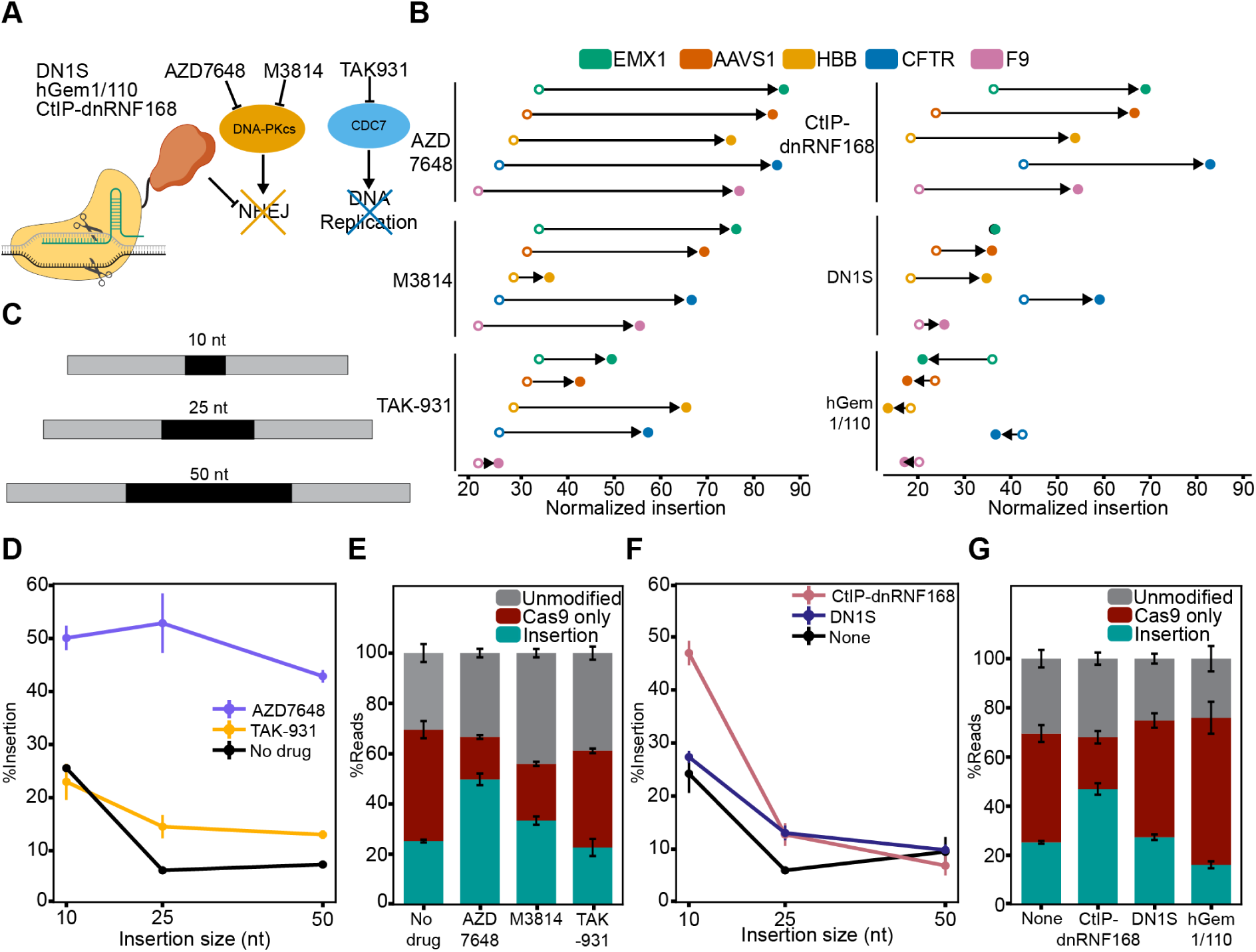
Inhibiting non-homologous end joining boosts templated insertion. A) Two strategies that boost templated insertion. Left: Cas9 is fused to proteins that alter DNA repair pathway choice. Right: small molecule inhibition of DNA-dependent protein-kinase catalytic subunit (DNA-PKcs) or CDC7. B) The effect of inhibitors (left) and Cas9 fusions (right) on the relative rate of templated insertion at five loci. AZD7648 (left, top) and Cas9-CtIP-dnRNF168 (right, top) both have the strongest effect at all tested loci. Open circles: editing with no inhibitor or DNA repair protein. Closed circle: editing with the indicated inhibitor or DNA repair protein fused to Cas9. All circles indicate the mean across three biological replicates. Arrow: change in editing efficiency with the indicated inhibitor. Fusions increase the relative rates of templated repair across all loci. Closed and open circles: mean of three biological replicates. C) Schematic of experiments with 50 nt homology arms and increasing insert lengths at the EMX1 locus. D) AZD7648 increases the insertion efficiency across all cargo sizes tested in this study. Error bars: mean of three biological replicates. E) AZD7648 outperforms TAK-931 and M3814 in boosting 10 nt insertion efficiency at EMX1 without increasing mutational signature or Cas9-generated indels. Error bars: mean of three biological replicates. F) Insertion efficiency decreases for all Cas9-repair protein fusions at EMX1. Error bars: mean of three biological replicates. G) Cas9 fused to CtIP-dnRNF168 increased insertion efficiency of 10 nt cargo at EMX1 without increasing mutational signature or Cas9-generated indels compared to no DNA repair fusion, DN1S, and hGem1/110. Error bars: mean of three biological replicates.

Cas9 fusion proteins can improve templated DNA insertion locally without perturbing repair pathways globally [53]. We focused on three fusions that have been previously characterized across multiple loci and cell types (**Figs. 4A, B, right**) [73–75]. Fusing Cas9 with the HDR-promoting CtIP and a dominant negative RNF168 (dnRNF168) increased retron editing efficiency 1.8-2.5 fold across five loci. A dominant-negative mutant of 53BP1 (DN1S) had variable effects across the five loci, with no improvements at EMX1. (**Fig. 4B, right**) [74]. Fusion of Cas9 to the N-terminal region of human Geminin limits Cas9 expression to the S/G2/M phases of the cell cycle when HDR is most active [76]. However, Cas9-hGem1/100 fusions either decreased or had no effect on retron editing activity (**Fig. 4B, right**). This result may reflect the mechanistic differences between ssODN and double-stranded donor DNA repair pathways (see **Discussion**). Combining Cas9-CtIP-dnRNF168 fusions with AZD7648 and TAK-931 did not improve editing any further at EMX1 (**Fig. S5B**). This finding is consistent with the hypothesis that Cas9-CtIP-dnRNF168 and AZD7648 both down-regulate NHEJ, albeit via different mechanisms. In contrast, retron-mediated insertion with Cas9-DN1S and Cas9-hGem1/110 fusions was higher with AZD7648 and TAK-931, but never exceeded AZD7648 or Cas9-CtIP-dnRNF168 alone. We conclude that selective inhibition of DNA-PKcs significantly improves retron-mediated gene editing.

Next, we tested retron editing activity with the nickase Cas9(D10A), and its fusions with DNA repair proteins (**Fig. S5C-E**). On its own, Cas9(D10A) paired with Efe1-RT caused an insertion in 1% of all reads at EMX1. However, fusing Cas9(D10A) to CtIP-dnRNF168 and a hyperactive HR recombinase hRad51(K133R) increased insertion rates >20-fold [77] (**Fig. S5D**). We also reasoned that fusing Cas9(D10A) to a helicase could improve HDR by peeling back the target and non-target DNA strands for the retron-synthesized DNA. We tested this hypothesis with Rep-X, an engineered helicase with exceptional processivity [78]. Cas9(D10A)-Rep-X helicase fusions increased insertion efficiency >20-fold, suggesting that retron editors can be used for DSB-free editing. Additional studies will be required to further optimize Cas9(D10A)-retron-RT editors and their combinations with DNA repair factors (see **Discussion**).

### Retron editing in cell lines and vertebrates

As a proof of principle, we next used retron editors to insert a split GFP for live cell imaging of endogenously expressed proteins in U2OS cells. GFP1-10, comprised of the first 10 GFP *β*-strands, is expressed from an integrated & inducible promoter (**Fig. S6**). The 11^th^ *β*-strand is fused to the protein of interest via a short linker. GFP1-10 is not fluorescent until it is complemented by GFP11 because chromophore maturation requires a critical GFP11-encoded glutamic acid [79]. Thus, fusing GFP11 to a target protein allows visualization of sub-cellular localization in live cells [80–82].

We targeted two proteins for endogenous GFP11 tagging via retron editors. The dispensable msd region was replaced with a 197 nt DNA template that included 70 nt homology arms, a nine nt linker encoding Gly-Gly-Gly, and the 48 nt GFP11 epitope. To maximize the integration efficiency, cells were transfected with Cas9-CtIP-dnRNF168 (see **Channeling repair pathway choice increases insertion efficiency**). 72-hours post-transfection, GFP1-10 was induced for 48 hours and GFP+ cells were collected using fluorescence-activated cell sorting (FACS). Confocal imaging of the insertion region confirmed the genomic edit (**Fig. S6C**). These results demonstrate that Efe1-RT can synthesize 200 nt ssDNAs. More broadly, this approach can be readily used for installing epitopes, disease-specific alleles, and other large insertions across the entire proteome from a genetically encoded cassette.

To expand retron editors beyond plasmid-based expression, we tested delivery in an all-RNA format (**Fig. 5**). In these experiments, Cas9 and Efe1-RT mRNAs were *in vitro* transcribed, 5’-capped, and 3’-polyA-tailed. The msr-msd was *in vitro* transcribed, and the sgRNAs were chemically synthesized (**Fig. 5A**). The four RNA cocktail were transfected into HEK293T cells and editing efficiency was scored by deep sequencing. Insertion of a 10 nt cargo was 9.9, 11.4, and 9.3% for the *EMX1*, *CFTR*, and *AAVS1* loci, respectively (**Fig. 5B**). This lower editing efficiency is likely due to the reduced Cas9 and RT expression from mRNA relative to plasmid-based delivery. Optimizing the UTRs, chemical modifications, and mRNA to ncRNA ratio may further boost editing at therapeutically relevant loci. We conclude that retron editors are compatible with direct RNA delivery.

**Figure 5:**
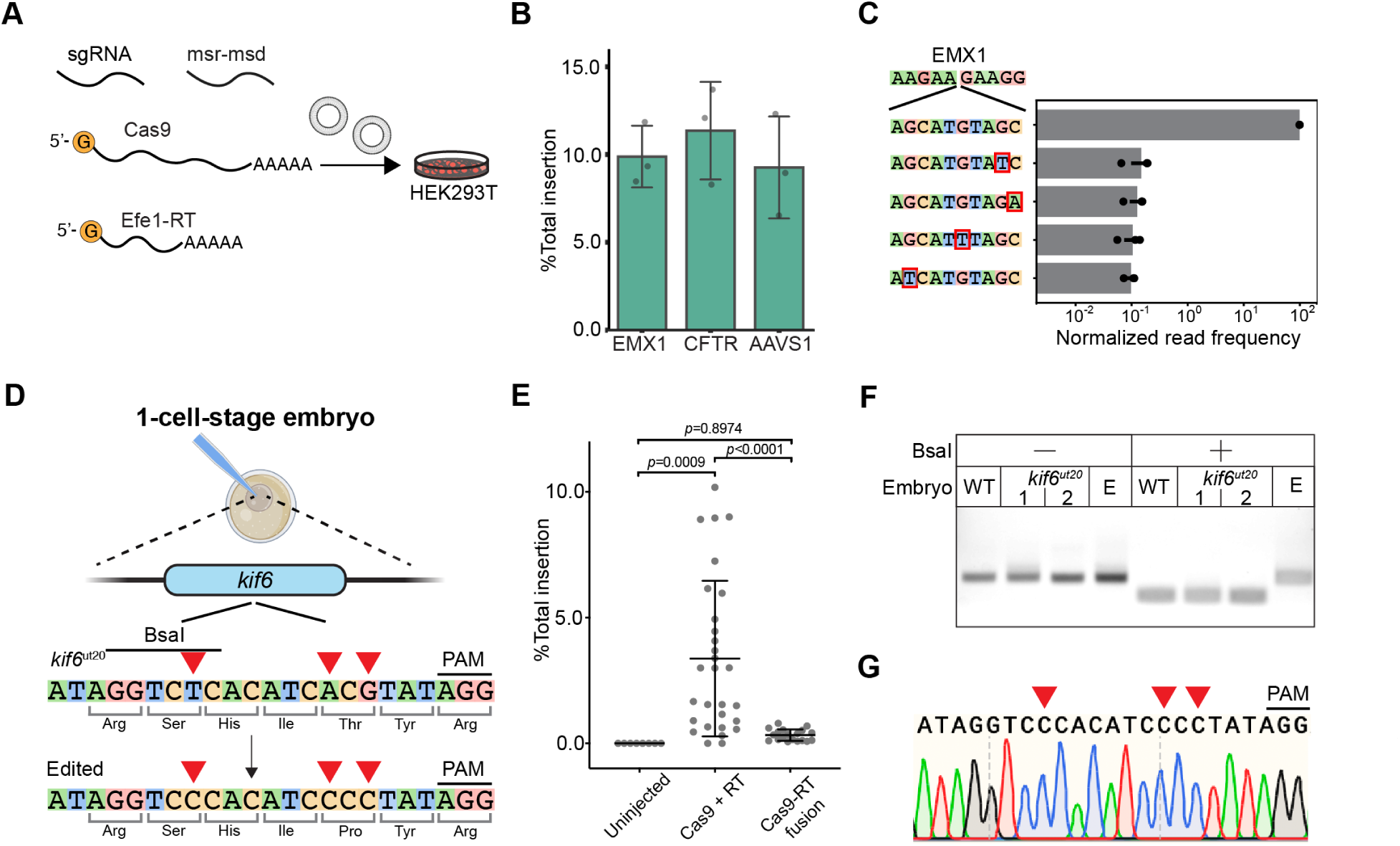
Retron editor delivery via an all-RNA package. A) Illustration of RNA-based retron editing. Cas9 and Efe1-RT are delivered as capped and poly-A-tailed mRNAs. The sgRNA and msr-msd are mixed with the mRNAs in a 10:1 molar ratio before transfection into HEK293T cells. B) Insertion efficiency of a 10 nt insert at the indicated loci with RNA-based delivery. Error bars: mean of three biological replicates (dots). C) The relative insertion frequency of a 10 nt cargo at the EMX1 locus, along with the four most frequent misincorporated sequences. Error bars: mean of three biological replicates (dots). D) Schematic of the three substitutions that are introduced in *kif6* by mRNA injection. The first mutation is silent but abolishes a BsaI cut site. E) Deep sequencing of embryos 24 hr post RNA injection for the indicated conditions. Each point is a single embryo. p-values are determined by a One-Way ANOVA test. F) BsaI restriction enzyme digests of the edit site sub-cloned from WT, *kif6*Δ, and edited embryos. The BsaI cut site is abolished in edited, but not WT or*kif6*Δ embryos. G) Sanger sequencing of the edited embryo from (E) confirms precise editing at the three expected sites (red triangles).

We next tested the repair of a pathogenic mutation via mRNA-based retron editing in zebrafish. We designed a sgRNA to target a pathogenic mutation in the Kinesin Family Member 6 *kif6^ut^*^20^ gene. Mutations in *kif6^ut^*^20^ cause scoliosis in zebrafish and are linked to neurological defects in humans [83]. The msd was designed to correct two base mutations that reverted a pathogenic Pro*→*Thr substitution. In addition, we introduced a silent T*→*C mutation that abolished a BsaI cleavage site for downstream restriction enzyme analysis (**Fig. 5E-F**). Embryos were injected with the sgRNA, msr-msd, and a fused Cas9-RT or split Cas9 and Efe1-RT mRNAs. Genomic DNA was harvested 24 hours post-injection, and submitted to NGS, restriction enzyme digestion, and Sanger sequencing (**Fig. 5D-F**). As expected, untreated controls did not show any editing. Injecting Cas9 and Efe1-RT mRNAs, along with the two ncRNAs, induced edits up to 10% of all reads from crude genomic preparations. The fused Cas9-RT mRNA showed reduced editing as compared to the split mRNAs (**Fig. 5E**). This may be due to the stability of the fusion mRNA in zebrafish. To further confirm retron editing, we sub-cloned the genomic DNA and subjected it to restriction enzyme and Sanger sequencing analysis (**Fig. 5F-G**). BsaI treatment of the edited, but not WT and *kif6^ut^*^20^ embryos resulted in a single band, indicating the expected cleavage pattern. Sanger sequencing of the insert site sub-cloned from an edited embryo also showed the expected edits. Collectively, these results indicate that retron editors are active in cell lines and zebrafish embryos when delivered as an all-RNA package.

## Discussion

Here, we engineer retron editors for precise genome engineering in mammalian cells and zebrafish embryos. Our metagenomic survey yielded 17 RTs that outperform Eco1-RT in mammalian cells. Efe1-RT, the lead candidate, is highly active, specific for its cognate RNA, and is capable of generating at least *∼*200 nt ssDNAs *in vivo* (**Fig. 1**). The most active RTs in our survey were all derived from clade 9. A recent bacterial functional screen also concluded that RTs in this clade generate high ssDNA levels in *E. coli* [17]. Additional structure-function studies will be required to dissect the mechanistic basis for this higher activity. Finally, we demonstrated retron editor delivery in an all-RNA format (**Fig. 5**). We anticipate that further optimizations, such as increasing mRNA and ncRNA stability will further improve editing efficiency (see below).

We iteratively improved on prior designs by optimizing the NLS, nuclease-RT linkers, and nuclease combinations (**Fig. 3**). The linker and NLS are key design components for both base and prime editors [84, 85]. In contrast, we show that retron editors can accommodate diverse NLS and linker combinations. Importantly, retron editors are compatible with Cas9, Cas12a, and even the nickase Cas9(D10A) (**Fig. 3, S5**). Cas12a-based retron editors further expand the potential target range and create opportunities for multiplexed retron editing due to Cas12a’s ability to process its crRNA [84]. Based on our results, we anticipate that retron-RTs will be compatible with other established, i.e. transcription activator-like effectors (TALEs), and emerging RNA/DNA-guided nucleases. Importantly, further development of nickase-based retron editors promises to avoid the induction of double-stranded DNA breaks (**Fig. S5**).

Retron editors are uniquely capable of synthesizing high copy numbers of the repair template at the excision site [48, 86, 87]. Boosting ssDNA synthesis via RT and ncRNA engineering will further improve the processivity, fidelity, and ultimately, edit length and efficiency. For example, rational engineering of a retron RT-based prime editor boosted editing efficiency >8-fold [11]. The design of the msr-msd and repair template relies on empirical testing of different msr-msd designs and offers an additional area for improvement. This includes modifying the insertion and truncation site of the native msd, homology arm length, target/non-target strand selection, and overall RNA structure. RNA circularization, structured RNA pseudoknots, and chemical modifications also increase ncRNA stability in mammalian cells [88]. Further optimization in these directions may enhance the overall efficiency of retron editing. In general principles and predictive algorithms for msr-msd and repair templates will further improve retron editors.

The repair of DSBs with a single-stranded DNA donor proceeds via single-strand template repair (SSTR) [89–91]. Our results are also consistent with an SSTR-based repair mechanism for retron editors. First, NHEJ inhibitors increase retron editing and reduce NHEJ-associated indels for both Cas9 and Cas12a across all tested target sites (**Fig. 4**). Second, fusing Cas9 with a dominant-negative allele of 53BP1, or with the DNA resection promoting CtIP, also increases retron editor efficiency. Third, we observed that 36-50 nt homology arms are optimal for retron editors (**Fig. 2**). Similarly, SSTR is maximized with 30-60 nt homology arms [92, 93]. SSTR is a RAD52-dependent process in yeast and human cells, suggesting that nuclease-RAD52 and/or RT-RAD52 fusions may boost retron editing [75, 94]. SSTR competes with two error-prone DSB repair pathways: classical NHEJ and polymerase theta-mediated end joining (TMEJ) [52, 95]. Dual inhibition of NHEJ and TMEJ may further synergize with RAD52 fusions [47, 69]. Rational design of asymmetric templates, cleavage-blocking mutations, and dual Cas9 nickases are additional avenues for maximizing editing efficiency [93, 96]. Mechanistic studies of retron editor-mediated repair will further improve editing outcomes across all domains of life.

In conclusion, retron editors are emerging as a highly promising gene editing tool. Their unique ability to accurately insert or replace sizeable DNA segments opens up possibilities for correcting complex genetic mutations that were previously challenging to address. Compatibility with an all-RNA formulation opens new avenues for therapeutic delivery into cells and organisms. Additionally, retron editors are poised to broaden the scope of high-throughput functional screens, allowing for the characterization of complex genetic variants with single-base resolution. Integration of retron editors into existing screening pipelines holds great promise for advancing our understanding of gene function and regulation, ultimately paving the way for novel therapeutic interventions and biotechnological applications.

## Materials and Methods

### Oligonucleotides and Plasmids

Retron-RT and ncRNAs gene blocks were ordered from IDT or Twist Biosciences and cloned into a GFP dropout entry vector via Golden Gate assembly. mRNAs were purchased from Cisterna Biologics. Retron editor expression plasmids were assembled by combining the sgRNA, retron msr-msd, the RT, and SpCas9 or AsCas12a in a *ccdB* dropout mammalian expression vector using Golden Gate assembly.

### Tissue Culture

HEK293T cells were generously provided by Professor Xiaolu A. Cambronne. HEK293T and U2OS were cultured in Dulbecco’s modified Eagle’s medium (DMEM) with 10% fetal bovine serum (Gibco) and 1% Penicillin-Streptomycin (Gibco). All cell lines were cultured and maintained at 37 °C and 5% CO_2_. U2OS Flp-In TREx - GFP1-10 were generated by stably integrating a GFP1-10 construct at the single FRT locus through dual transfection of pcDNA5/FRT/TO/Intron-eGFP1-10 and pOG44 plasmids in a 1:10 ratio and subsequent selection with hygromycin at 200 µg/mL and blasticidin at 15 µg/mL.

### Metagenomic Discovery

To find new retrons, we searched the NCBI genome and metagenomic contigs for the retron-RT gene using hidden Markov models (HMMER, [97, 98]) and a database containing all experimentally validated retron-RT with a permissive e-value threshold of 10*^−^*^4^ [33, 49, 50]. We next established a pipeline to detect structured RNAs near retron RT sequences. Initially, each experimentally validated msr-msd transcript and its close counterparts served as seeds for CMfinder 0.4.1, an RNA motif predictor that leverages both folding energy and sequence covariation [99]. Covariate models were then crafted with Infernal suite’s cmbuild and used to search for analogous structures around the start of the RT open reading frame (ORF). We manually inspected the msr-msd regions of all retron candidates, especially those that didn’t return any hits via the automated pipeline. RNAfold was used to inspect structured regions, and to compare them to known msr-msd transcripts [100]. We then used MAFFT-Q-INS-i for multiple alignments, focusing on identifying conserved sequences in related genomes. Subsequently, we pruned the alignments at the a1 and a2 areas, cycling back to earlier pipeline steps to acquire more sequences from both ends. R-scape was used to pinpoint covarying base pairs in the suggested consensus structures, mitigating the influence of phylogenetic correlations and base composition biases not attributed to conserved RNA structure. Accessory retron genes adjacent to the RT were annotated using the Pfam database as a query and HMMER. High-confidence retron systems were manually inspected to confirm the expected protein domains, catalytic residues, and ncRNA structure. To remove redundant sequences, all putative hits were clustered with CD-HIT [101], setting a sequence identity threshold at 90% and an alignment overlap of 80%. The clustered datasets were then transformed into phylogenetic trees, and candidates for validation were selected based on their positioning in the tree.

To organize the retron RTs into phylogenetic groups, we employed MAFFT software and used progressive methods for multiple sequence alignments (MSAs) [102]. An MSA was constructed from the RT0–7 domain of 1,912 sequences, sourced from a dataset of 9,141 entries previously categorized as retron/retron-like RTs and an additional 16 RTs from experimentally verified retrons [33]. Phylogenetic trees were generated using FastTree, applying the WAG evolutionary model, combined with a discrete gamma model featuring 20 rate categories. Specifically, the RT tree was crafted using IQ-TREE v1.6.12, incorporating 1000 ultra-fast bootstraps (UFBoot) and the SH-like approximate likelihood ratio test (SH-aLRT) with 1000 iterations [103]. The best-fit model, identified by Modelfinder as the LG+F+R10 due to its minimal Bayesian Information Criterion (BIC) value among 546 protein models, was used. The RT Clades’ internal nodes exhibited UFBoot and SH-aLRT support values exceeding 85% [101].

### Plasmid-based fluorescent reporter assays

1.2*×*10^5^ HEK293T cells were seeded in a 24-well plate 18-24 hours before transfection. 0.35 µg of the retron editor plasmid and 0.35 µg of the fluorescent reporter plasmid were co-transfected using Lipofectamine 2000 (Invitrogen). Cells were trypsinized and collected for flow analysis 72 hours after transfection. Flow analysis was conducted on a Novocyte flow cytometer (ACEA Biosciences). Cells were gated to exclude dead cells and doublet and 10,000 cells were analyzed in all samples. Cells were then gated by FITC-A (x-axis) and PE-Texas Red-A (y-axis). The editing efficiency was reported as the percentage of cells in the quadrant of FITC-A and PE-Texas Red-A.

### Genomic reporter assays

The fluorescent reporter was cloned in a plasmid designed for Bxb1 recombinase-driven landing pad system [104]. This plasmid was transfected into landing pad HEK293T cells followed by doxycycline induction and AP1903 selection to generate stably integrated reporter cells. 1.2 *×* 10^5^ of HEK293T reporter cells were seeded in a 24-well plate 18-24 hours before transfection. 1 µg of the retron editor plasmid was transfected using Lipofectamine 2000. 48-72 hours after transfection, cells were treated with doxycycline to induce the expression of the fluorescent reporter. Cells were trypsinized and collected for flow analysis and genomic DNA extraction (Qiagen DNeasy Blood and Tissue kit).

### Confocal imaging

HEK293T cells were seeded and transfected as described for the plasmid-based fluorescent reporter assay (see above). 48 hours post-transfection, cells were seeded into 15 mm glass-bottom cell culture dishes (NEST) and incubated for an additional 24 hours. Cells were then imaged with a Nikon Ti2 Spinning Disk Confocal Microscope at 20x magnification. 8858 x 8858-pixel images were acquired and processed using ImageJ.

U2OS FlpIn TREx GFP1-10 cells were transfected with 1 µg retron editor plasmids that target either the N- or C-terminus of the protein of interest to insert a GFP_11_ fragment. 48-72 hours after transfection, cells were expanded into 10-cm plates. Cells with high GFP intensities were then sorted via a cell sorter (Sony MA900). For confocal imaging, cells were incubated with doxycycline for 48-72 hours before imaging to induce the expression of the GFP1-10 construct. Image acquisition was performed with live cells under spinning-disk confocal microscopy (Olympus).

### Next generation DNA sequencing

Genomic samples were subjected to two rounds of PCR for NGS library preparation 2. The first round of PCR was performed using the KOD One PCR master mix (TOYOBO). Primers were designed *∼*600 base pairs away from the cut site on each side to avoid amplifying the RT-generated ssDNA. PCR products were gel purified and barcoded via a second round of PCR with Illumina P5/P7 adapters using Q5 HotStart High Fidelity master mix (NEB). PCR amplicons were sequenced on an Illumina Novaseq. Reads were demultiplexed using NovaSeq Reporter (Illumina). Alignment of amplicon sequences to a reference sequence was performed using CRISPResso2 [105]. The aligned sequences were first checked for the expected cargo insertion. Next, the homology arms and insertion sequences were further checked for any mismatches or deletions. We considered a “perfect edit” as any sequence that had the expected insertion only, with no additional mismatches and deletions. If these events also had a mismatch or deletion, we counted it as an “imperfect” edit. Any insertions, deletions, or mismatches that did not have the intended cargo were surmised to be from nuclease-only activity.

### In vitro transcription of the msr-msd

The msr-msd was PCR amplified from a plasmid or gene block with a T7 promoter. The ncRNA was generated using the HiScribe T7 High Yield RNA Synthesis Kit (NEB) according to the manufacturer’s protocol. RNA products were purified using the RNeasy Mini Kit (Qiagen).

### Zebrafish Maintenance and Gene Editing

All zebrafish experiments were performed according to University of Texas at Austin IACUC standards. Embryos were raised at 28.5 °C in fish water (0.15% Instant Ocean in reverse osmosis water). Wildtype and *kif6*^ut20(P293T)^ mutants were in-crossed, and one-cell stage embryos were injected with 1 nL of the injection mix using a microinjector-pump system (World Precision Instruments Nanoliter Injector and PV 820 Pneumatic PicoPump). Injected embryos and un-injected sibling controls were incubated at 28.5 °C in fish water until 24 hr post fertilization, at which point, surviving embryos were euthanized in excess Tricaine (0.4% MS-222). Genomic DNA was extracted from individual embryos using the HotSHOT Method [106]. Briefly, embryos were transferred into 50 mM NaOH and heated to 95 °C for 20 min. The samples were neutralized with ^1^ volume of 1 M Tris-HCl, pH 8, before downstream analysis. For NGS sequencing, genomic DNA was PCR amplified to extend the amplicon with Illumina adapters using the KOD One PCR master mix (TOYOBO). PCR amplicons were directly sequenced on an Illumina NovaSeq sequencer.

## Data Availability

Raw FASTQ files and processed data tables are available at GenBank. Source data are provided with this paper.

## Declarations

### Author Contributions

K.J., H.-C.K., K.H. and I.J.F. conceived the project. K.H. carried out the bioinofrmatic discovery and NGS data analysis. J.B., H.-C.K., Y.-C.C. and K.J. performed all cell line experiments and analyzed the data. B.V. and R.S.G. performed the zebrafish experiments. Y.-R.L. S.K.D., and B.X. established the U2OS cell lines and performed some experiments. M.E.L. assisted with cloning. J.B., H.-C.K., K.H., Y.-C.C., and I.J.F. prepared the figures and wrote the manuscript with input from all co-authors. I.J.F. secured the funding and supervised the project.

## Acknowledgments

We thank Daphne Sahaya for cloning assistance, and all members of the Finkelstein lab for discussions. Professor Xiaolu A. Cambronne provided access to a flow cytometer and the analysis software. Confocal imaging was conducted at the Center for Biomedical Research Support with the assistance of Paul Oliphint. This work was supported by a Sponsored Research Agreement from Retronix (to I.J.F.), the Welch Foundation grant F-1808 (to I.J.F.), and the College of Natural Sciences Catalyst grant for seed funding.

