## Supplementary Information for "Discovery and Engineering of Retrons for Precise Genome Editing"

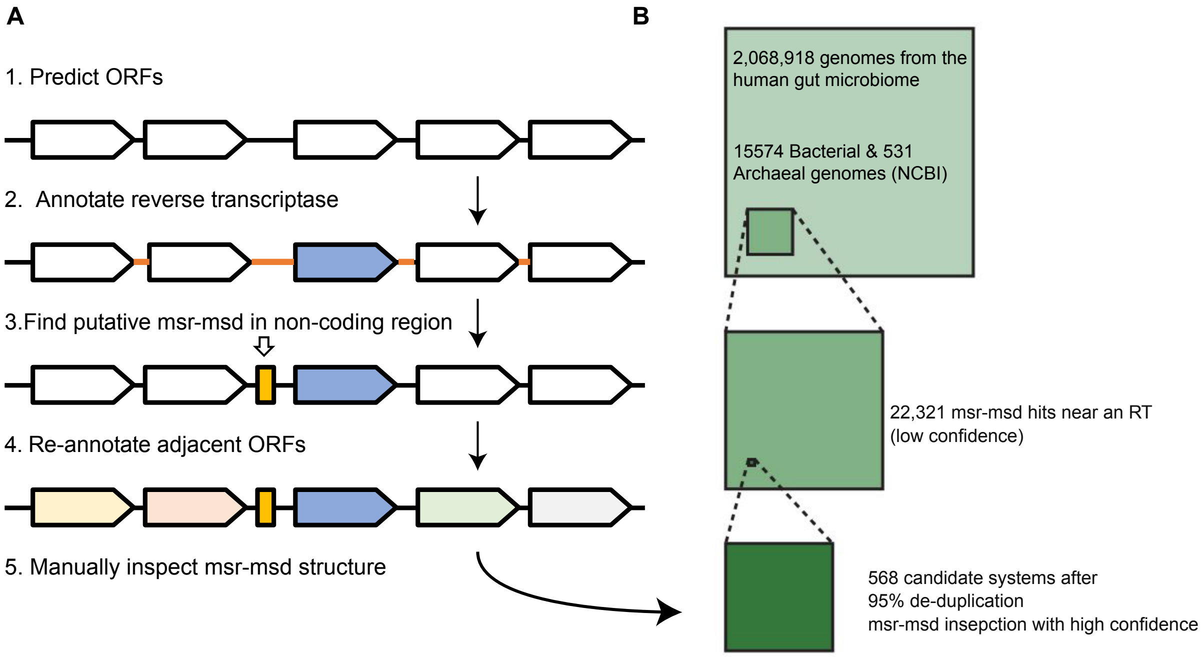


**Figure S1: Overview of the bioinformatic retron discovery pipeline.** A) The pipeline involves five steps: 1. Predict open reading frames (ORFs) with Prodigal [1]; 2. Annotate reverse transcriptase (RT) genes using HMMER [2]; 3. Identify putative msr-msd sequences in non-coding regions using cmfinder and infernal [3, 4]; 4. Re-annotate adjacent ORFs with HMMER; and 5. Manually inspect msr-msd structures with ViennaRNA. B) The analysis was conducted on 2,068,918 reference genomes from the human gut microbiome [5], along with 15,574 bacterial and 531 archaeal genomes from the NCBI database [6]. After a $95\%$ de-duplication at the amino acid sequence level and annotating msr-msds, we identified 568 new candidate systems.

‘

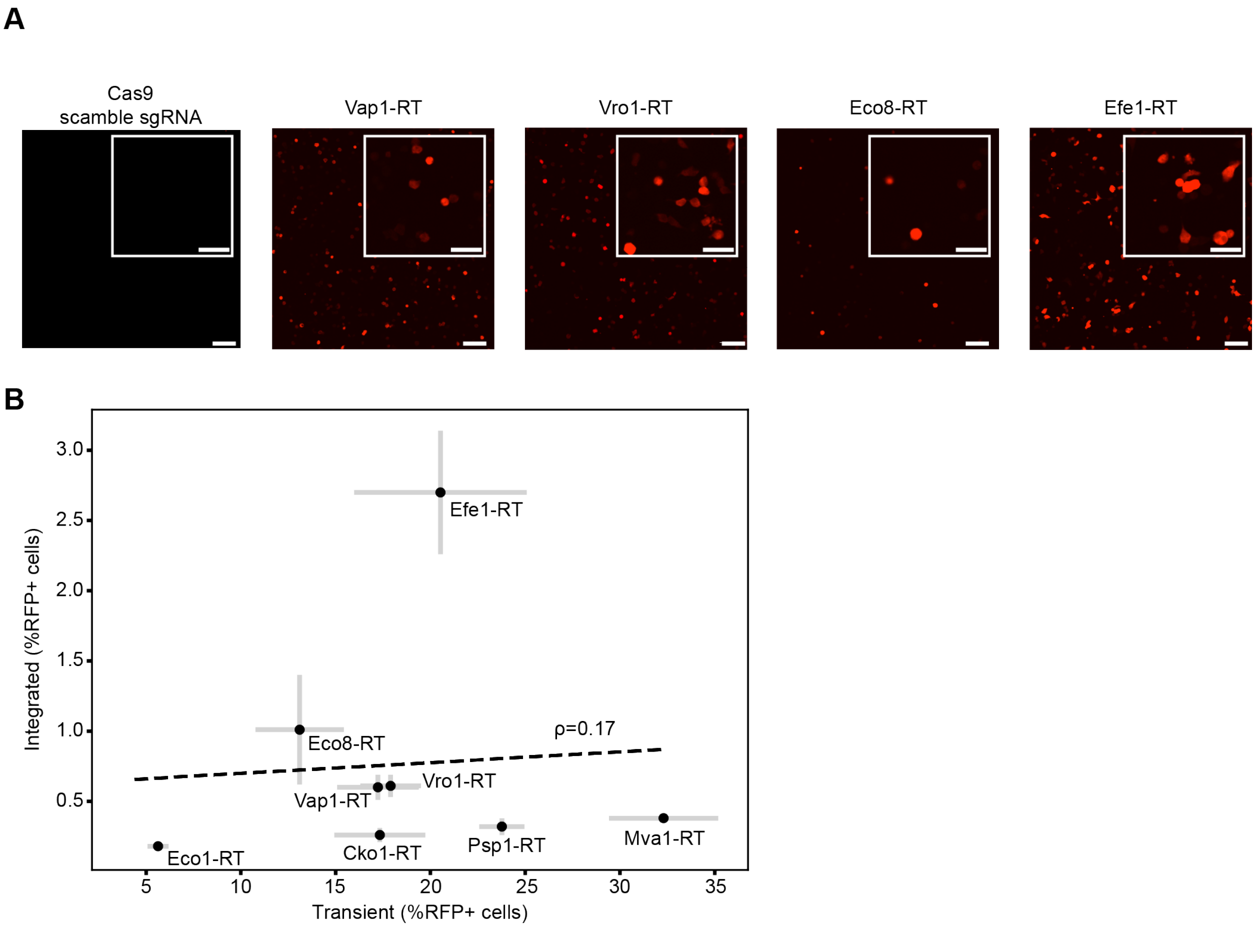


**Figure S2: Comparison of top RTs in transient and genomically-integrated RFP reporter assays.** A) Confocal images of the indicated RTs, or Cas9 with a scrambled sgRNA. Scale bar: 100 µm; inset: 50 µm. B) Correlation between retron editing activity in a genomic vs. transient RFP reporter system. The genomic RFP reporter was integrated into the AAVS1 locus. Error bars: mean of three biological replicates.

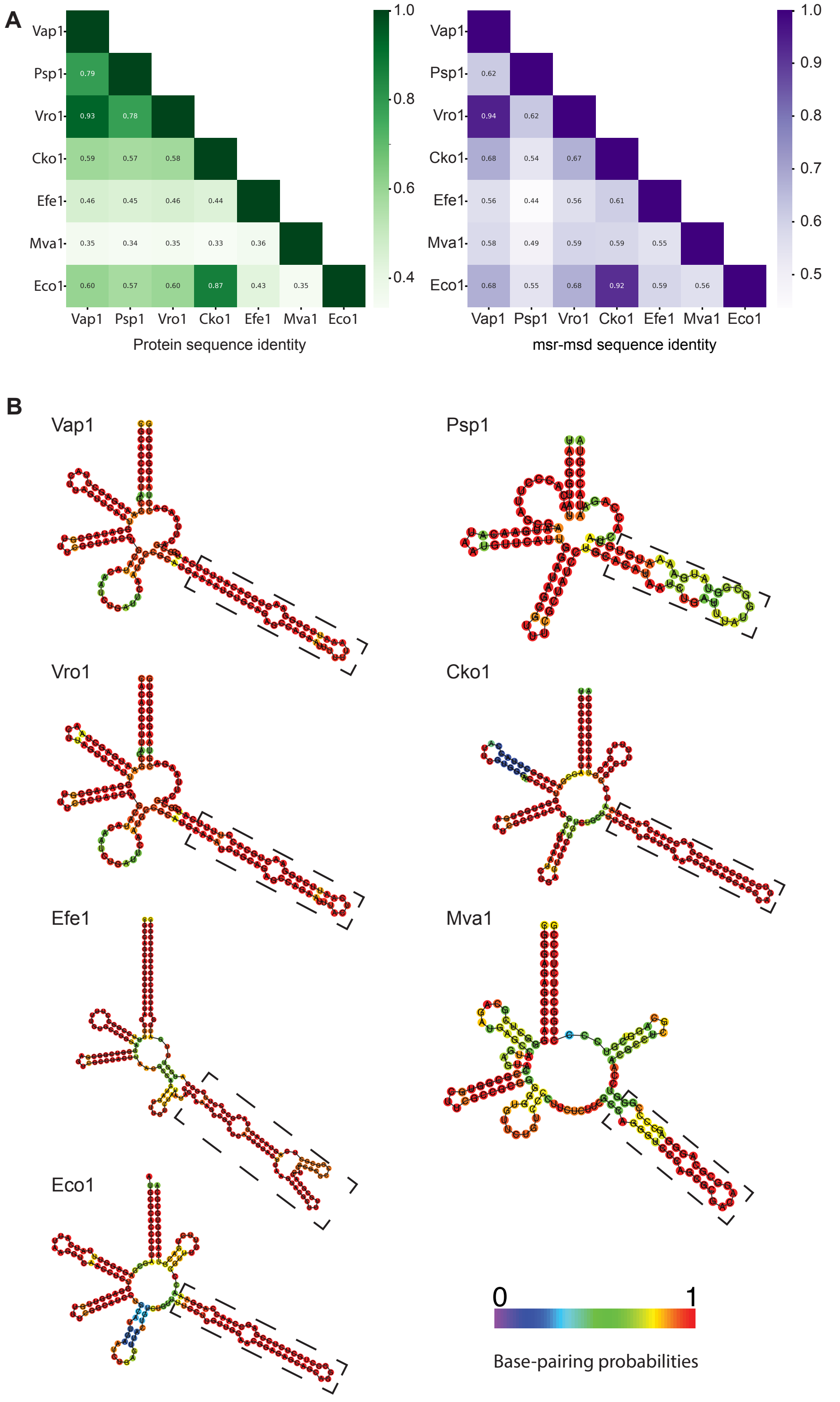


**Figure S3: Sequence identity and structural analysis of highly active retron-RTs.** A) Sequence identity heatmap of RT's amino acid sequences and ncRNA sequences for retron candidates. B) Structure of representative msr-msd transcripts encoded by retron candidates. The structures are predicted by ViennaRNA. The msd region is highlighted by a dashed rectangle.

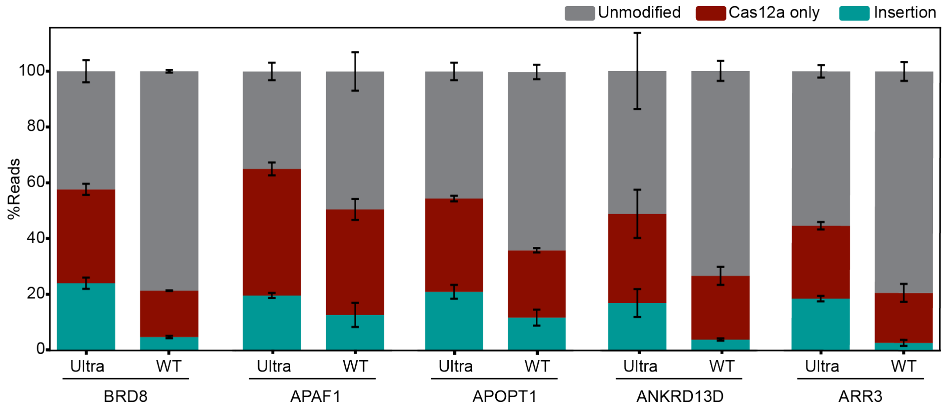


**Figure S4: AsCas12a-based editing outcomes at the indicated loci.** AsCas12a Ultra cleaves more efficiently at all loci that we tested in this study. Error bars: mean of three biological replicates.

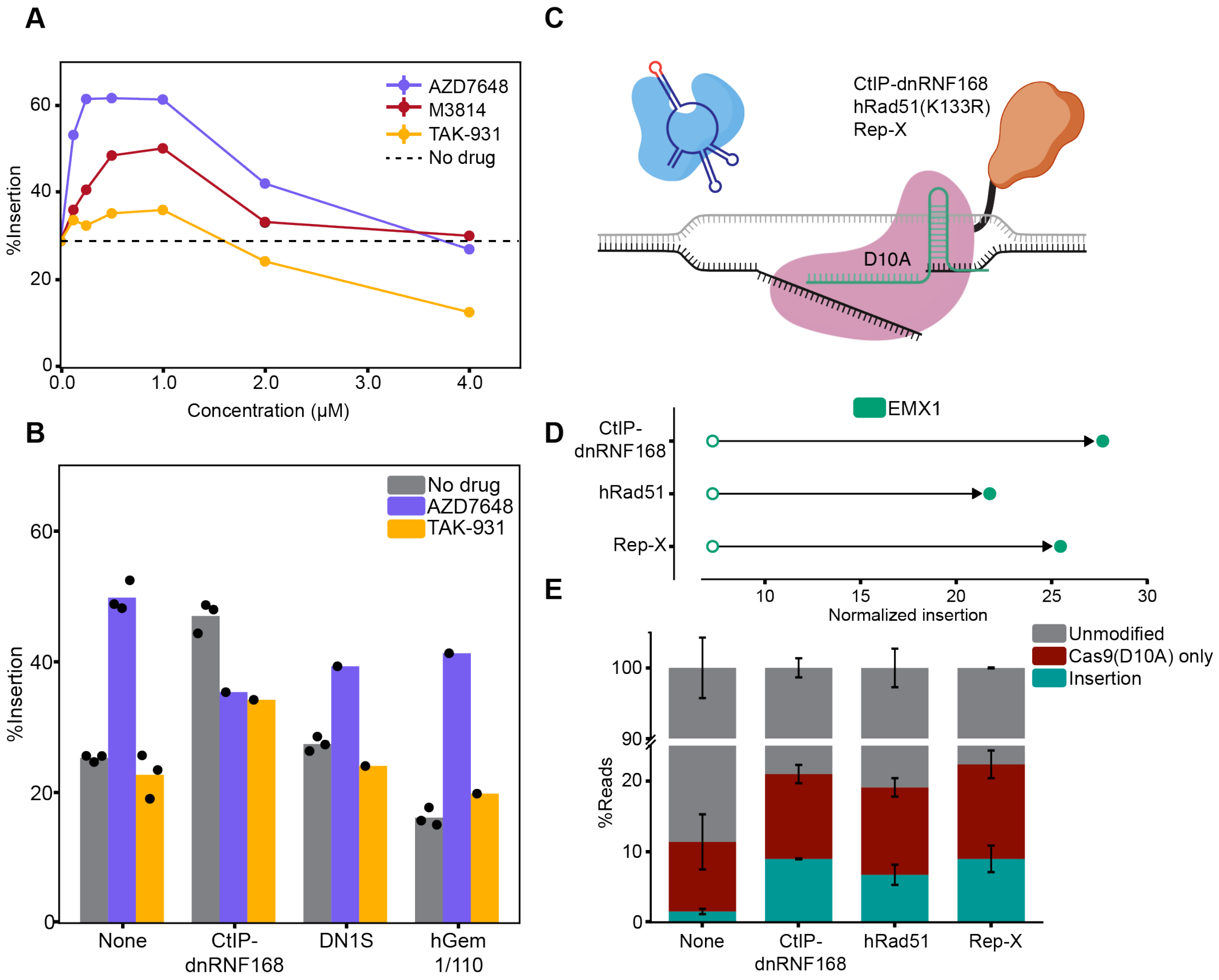


**Figure S5: Optimizing inhibitor and DNA repair fusions improves gene editing with nickase Cas9 (nCas9).** A) Optimization of inhibitor concentrations for optimal retron editing at the EMX1 locus. Dashed line: gene editing without inhibitors. B) The effect of combining DNA repair inhibitors with Cas9-DNA repair domain fusions. Combining the most active inhibitor, AZD7468, with Cas9-CtIP-dnRNF168 reduces overall insertion activity. In other cases, AZD7468 improves retron editing, likely due to the limited improvement observed with Cas9-DN1S and Cas9-hGem1/100 fusions. C) Illustration of the nickase Cas9(D10A)-DNA repair fusions, and a nickase-based retron editor. D) Fusing Cas9(D10A) with CtIP-dnRNF168, hRad51(K133R), and Rep-X helicase [7–9] increases the relative rates of templated repair at EMX1. Open circles: editing with no repair factor fusion. Closed circle: editing with the indicated repair factor fusion. All circles indicate the mean across three biological replicates. Arrow: change in editing efficiency with the indicated Cas9-DNA repair protein fusion. E) Breakdown of editing outcomes, as reported by NGS read counts. Cyan: retron editor-mediated insertion; Red: Cas9(D10A)-only edits without any insert, Gray: unmodified reads. Error bars: mean of three biological replicates.

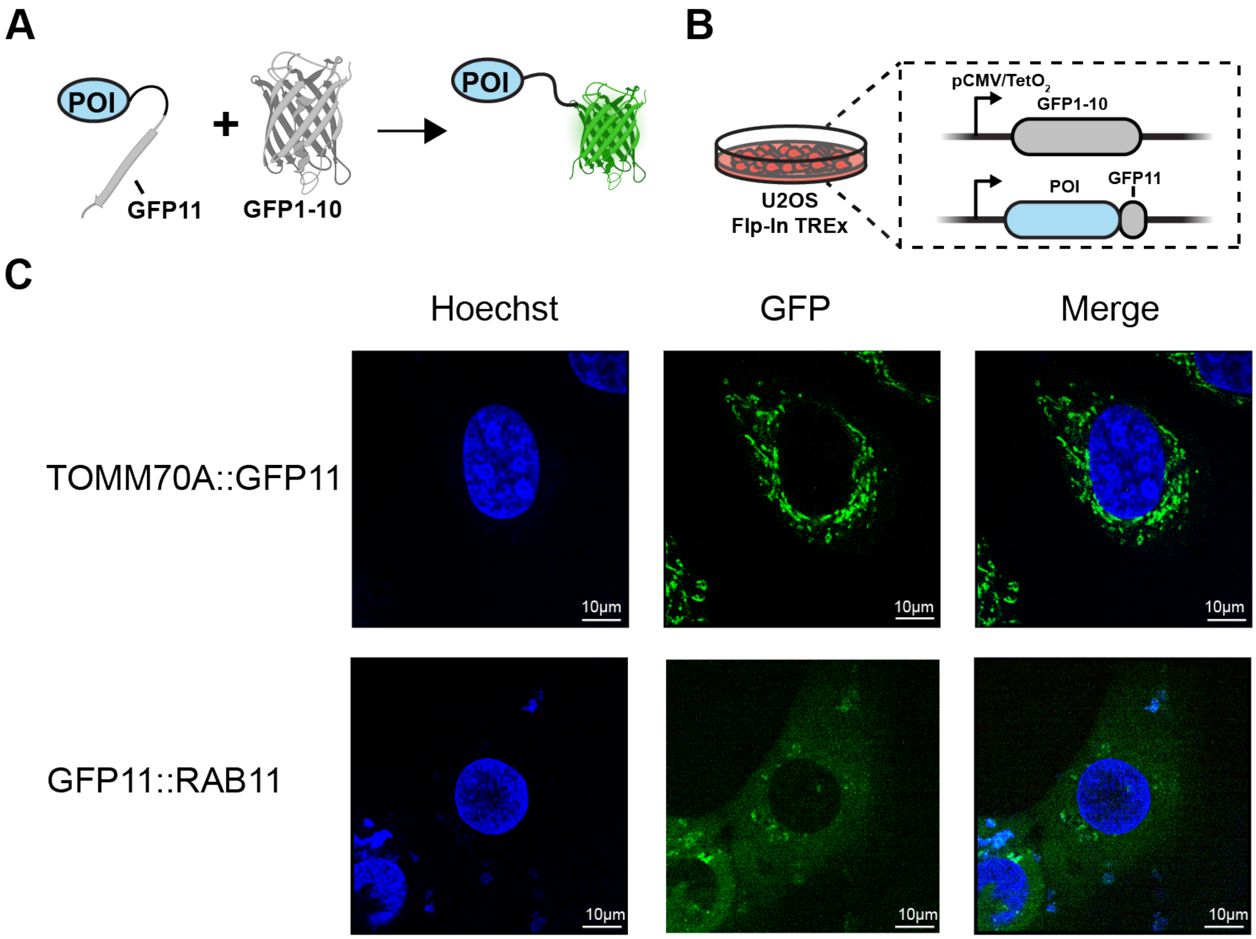


**Figure S6: In-frame precise epitope insertion with retron editors** A) Schematic of the split super folder GFP system. The 11th GFP $\beta$-strand (GFP11) is expressed as a fusion to the protein of interest (POI, blue). Reconstitution of GFP11 with GFP1-10 restores fluorescence. B) GFP1-10 is expressed from a genomically-integrated inducible promoter. GFP11 is inserted in-frame into the native genomic locus. C) Confocal images of two GFP11-protein fusions. Scale bars: 10 µm.

### Supplemental Tables

| Plasmid | Description | Source |
|---|---|---|
| pIF1056 | RFP(∆9)-GFP reporter | This paper |
| pIF1057 | Mammalian expression ccdB dropout | This paper |
| pIF1058 | sgRNA and msr-msd entry vectors | This paper |
| pI1F059 | SpCas9 entry vector | This paper |
| pIF1060 | Cas9(D10A) entry vector | This paper |
| pIF1061 | AsCas12a entry vector | This paper |
| pIF1062 | AsCas12a Ultra entry vector | This paper |
| pIF1063 | Efe1-RT entry vector | This paper |
| pIF1064 | Cex1-RT entry vector | This paper |
| pIF1065 | Eco8-RT entry vector | This paper |
| pIF1066 | Vap1-RT entry vector | This paper |
| pIF1067 | Vro1-RT entry vector | This paper |
| pIF1068 | SpCas9-CtIP-dnRNF168 entry vector | This paper |
| pIF1069 | SpCas9-DN1S entry vector | This paper |
| pIF1070 | hRad51(K133R) entry vector | This paper |
| pIF1071 | Rep-X entry vector | This paper |
| pIF1072 | SpCas9-SGGSx2-XTEN-SGGSx2-Efe1-RT sgRNA and msr-msd entry vector | This paper |
| pIF1073 | SpCas9 T2A Efe1-RT entry vector | This paper |
| pIF1074 | SpCas9 and Efe1-RT sgRNA and msr-msd entry vector | This paper |

**Table 1:** Plasmids used in this study.

| Name | Sequence, 5'→3' | Description |
|---|---|---|
| EMX1-outer-F | GACCCGCTTCCTCCCTGTCCTT | Outer PCR for EMX1 |
| EMX1-outer-R | GTCTCAACCTGCCCAGCGGTTC | Outer PCR for EMX1 |
| EMX1-NGS-F | AATGATACGGCGACCACCGAGATCTACACTCTTTCCCTACACGACGCTCTTCCGATCTGGCCTCCTGAGTTTCTCATC | NGS primer for EMX1 |
| EMX1-NGS-R | CAAGCAGAAGACGGCATACGAGAT[BC]GTGACTGGAGTTCAGACGTGTGCTCTTCCGATCTCTCTGCCCTCGTGGG | NGS primer for EMX1 |
| AAVS1-outer-F | GTTTCTTAGGATGGCCTTCTCC | Outer PCR for AAVS1 |
| AAVS1-outer-R | TCTGGGCGGAGGAATATGT | Outer PCR for AAVS1 |
| AAVS1-NGS-F | AATGATACGGCGACCACCGAGATCTACACTCTTTCCCTACACGACGCTCTTCCGATCTGAAATGGGGGTGTGTCACC | NGS primer for AAVS1 |
| AAVS1-NGS-R | CAAGCAGAAGACGGCATACGAGAT[BC]GTGACTGGAGTTCAGACGTGTGCTCTTCCGATCTGCTGCCTCCAGGGATC | NGS primer for AAVS1 |
| CFTR-outer-F | GTTTACTAGATTTCAGCCAGTTTTCAG | Outer PCR for CFTR |
| CFTR-outer-R | GATTAAACAAGTCTTGAATCTGTCTCC | Outer PCR for CFTR |
| CFTR-NGS-F | AATGATACGGCGACCACCGAGATCTACACTCTTTCCCTACACGACGCTCTTCCGATCTCTATGTTAAGGGAAATAGGACAAC | NGS primer for CFTR |
| CFTR-NGS-R | CAAGCAGAAGACGGCATACGAGAT[BC]GTGACTGGAGTTCAGACGTGTGCTCTTCCGATCTGTACAAATGAGATCCTTACCCC | NGS primer for CFTR |
| HBB-outer-F | TGTTCCCCCAGACACTCTTGCA | Outer PCR for HBB |
| HBB-outer-R | AGAATGGTGCAAAGAGGCATGA | Outer PCR for HBB |
| HBB-NGS-F | AATGATACGGCGACCACCGAGATCTACACTCTTTCCCTACACGACGCTCTTCCGATCTGCCAGGGCTGGGCATA | NGS primer for HBB |
| HBB-NGS-R | CAAGCAGAAGACGGCATACGAGAT[BC]GTGACTGGAGTTCAGACGTGTGCTCTTCCGATCTCACATGCCCAGTTTCTATTGG | NGS primer for HBB |
| F9-outer-F | GTGGAAGAGTTCTAAATGTGATCC | Outer PCR for F9 |
| F9-outer-R | GAAAAAACAAAGTGATATATGTTGCAT | Outer PCR for F9 |

Continued on next page

**Table 2:** (continued)

| Name | Sequence, 5'→3' | Description |
|---|---|---|
| F9-NGS-F | AATGATACGGCGACCACCGAGATCTACACTCTTTCCCTACACGACGCTCTTCCGATCTGTCAAATCATGTAATCAAAATTTAGTGAAG | NGS primer for F9 |
| F9-NGS-R | CAAGCAGAAGACGGCATACGAGAT[BC]GTGACTGGAGTTCAGACGTGTGCTCTTCCGATCTGTACTTTGGTACAACTAATCGACC | NGS primer for F9 |
| Innpl1a-NGS-F | AATGATACGGCGACCACCGAGATCTACACTCTTTCCCTACACGACGCTCTTCCGATCTCTTCCTGTCATGTGACCTTCAG | NGS primer for Innpl1a |
| Innpl1a-NGS-R | CAAGCAGAAGACGGCATACGAGAT[BC]GTGACTGGAGTTCAGACGTGTGCTCTTCCGATCTGTTTGCATGTGTGTGTGTTC | NGS primer for Innpl1a |
| ARR3-NGS-F | AATGATACGGCGACCACCGAGATCTACACTCTTTCCCTACACGACGCTCTTCCGATCTGGAGAGGTCTGTGGGT | NGS primer for ARR3 |
| ARR3-NGS-R | CAAGCAGAAGACGGCATACGAGAT[BC]GTGACTGGAGTTCAGACGTGTGCTCTTCCGATCTGTGGGGGTCACAAACC | NGS primer for ARR3 |
| APAF1-NGS-F | AATGATACGGCGACCACCGAGATCTACACTCTTTCCCTACACGACGCTCTTCCGATCTAGAATGCGGAGACGGTC | NGS primer for APAF1 |
| APAF1-NGS-R | CAAGCAGAAGACGGCATACGAGAT[BC]GTGACTGGAGTTCAGACGTGTGCTCTTCCGATCTCCATTTAAAGGAGTACTCCTGT | NGS primer for APAF1 |
| ANKRD13D-NGS-F | AATGATACGGCGACCACCGAGATCTACACTCTTTCCCTACACGACGCTCTTCCGATCTGCCCCTTGTCCTCTG | NGS primer for ANKRD13D |
| ANKRD13D-NGS-R | CAAGCAGAAGACGGCATACGAGAT[BC]GTGACTGGAGTTCAGACGTGTGCTCTTCCGATCTTCCCTGCCTTGCCC | NGS primer for ANKRD13D |
| BRD8-NGS-F | AATGATACGGCGACCACCGAGATCTACACTCTTTCCCTACACGACGCTCTTCCGATCTGGAGTCAAGTCCCCAAG | NGS primer for BRD8 |
| BRD8-NGS-R | CAAGCAGAAGACGGCATACGAGAT[BC]GTGACTGGAGTTCAGACGTGTGCTCTTCCGATCTTGCTTTAGTAATGCAACATACCT | NGS primer for BRD8 |

Continued on next page

**Table 2:** (continued)

| Name | Sequence, 5'→3' | Description |
| --- | --- | --- |
| APOPT1-NGS-F | AATGATACGGCGACCACCGAGATCTACACTCTTTCCCTACACGACGCTCTTCCGATCTGTTAATCCAATTTACTTTGTTAAAGG | NGS primer for APOPT1 |
| APOPT1-NGS-R | CAAGCAGAAGACGGCATACGAGAT[BC]GTGACTGGAGTTCAGACGTGTGCTCTTCCGATCTGTCAAATTCTGGTTTGCCCA | NGS primer for APOPT1 |
| KIF6_F | AATGATACGGCGACCACCGAGATCTACACTCTTTCCCTACACGACGCTCTTCCGATCTGCTACGTTTCCAGAATGAGCTT | Forward primer for genomic DNA amplification from the KIF6 locus for NGS |
| KIF6_R | CAAGCAGAAGACGGCATACGAGAT[BC]GTGACTGGAGTTCAGACGTGTGCTCTTCCGATCTTGTACACCGATGTTCTTCTTCTC | Reverse primer for genomic DNA amplification from the KIF6 locus for NGS |

**Table 2:** Oligonucleotides used to amplify genomic loci in this study. [BC] indicates a unique Illumina index.

| Locus | Spacer sequence, 5’→3’ | Type | Description |
|---|---|---|---|
| RFP | TGGCTACCAGCTTCATGCT | sgRNA | Targets RFP(Δ9) in the plasmid and integrated reporters |
| EMX1 | GAGTCCGAGCAGAAGAAGAA | sgRNA | Target EMX1 locus |
| AAVS1 | GGGGCCACTAGGGACAGGAT | sgRNA | Targets AAVS1 locus |
| CFTR | CTAAACTCATTAATGCCCTT | sgRNA | Targets CFTR locus |
| HBB | AGTCTGCCGTTACTGCCCTG | sgRNA | Targets HBB locus |
| F9 | CACTGAGTAGATATCCTAAA | sgRNA | Targets F9 locus |
| BRD8 | CTGCTTGACGAGCTAGAACAT | crRNA | Target BRD8 locus |
| APAF1 | ATCAGAAGCCCAGATTTGTCT | crRNA | Target APAF1 locus |
| APOPT1 | AGGTATGTAAAAGTGAACAGG | crRNA | Target APOPT1 locus |
| ANKRD13D | CCAGAGACACGGCCAGCTCCA | crRNA | Target ANKRD13D locus |
| ARR3 | CACCACCGGAGGCAGGCCCTG | crRNA | Target ARR3 locus |
| TOMM70A | GCACCAACATTATAAAACAG | sgRNA | Targets TOMM70A locus in U2OS |
| RAB11 | GGTAGTCGTACTCGTCGTCG | sgRNA | Targets RAB11 locus in U2OS |
| KIF6 | ATAGGTCTCACATCACGTAT | sgRNA | Targets Kif6 locus in zebrafish |

**Table 3:** Spacers for single guide RNAs (sgRNAs) and CRISPR RNAs (crRNAs) used in this study.

| Name | Amino acid sequence |
|---|---|
| C-myc | PAAKRVKLD |
| SV40 | PKKKRKV |
| BPSV40 | KRTADGSEFEPKKKRKV |
| vBPSV40 | KRTADGSE |
| NLP | AVKRPAATKKAGQAKKKKLD |
| 3xHA | YPYDVPDYAYPYDVPDYAYPYDVPDYA |
| Short NLS linker | EFE |

**Table 4:** Nuclear localization sequences used in this study.

| N-terminal NLS | C-terminal NLS | Rel. editing efficiency (mean ±S.D.) |
|---|---|---|
| SV40 | NLP | 1.00 ±0.09 |
| Cmyc-GG-BPSV40 | BPSV40-SV40 | 1.00 ±0.05 |
| Cmyc-GG-BPSV40 | BPSV40-SV40-EFE-NLP | 0.75 ±0.05 |
| Cmyc-GG-BPSV40 | BPSV40-vBPSV40-EFE-NLP | 0.76 ±0.11 |
| Cmyc-GG-BPSV40 | BPSV40-vBPSV40 | 0.73 ±0.02 |
| Cmyc-GG-BPSV40 | BPSV40-vBPSV40-EFE-BPSV40 | 0.70 ±0.07 |
| Cmyc-GG-BPSV40 | BPSV40-NLP-3xHA-SV40 | 0.49 ±0.02 |
| Cmyc-GG-vBPSV40 | BPSV40-SV40 | 0.97 ±0.08 |
| Cmyc-GG-vBPSV40 | BPSV40-SV40-EFE-NLP | 0.69 ±0.03 |
| Cmyc-GG-vBPSV40 | BPSV40-vBPSV40-EFE-NLP | 0.85 ±0.08 |
| Cmyc-GG-vBPSV40 | BPSV40-vBPSV40 | 0.82 ±0.04 |
| Cmyc-GG-vBPSV40 | BPSV40-vBPSV40-EFE-BPSV40 | 0.63 ±0.01 |
| Cmyc-GG-vBPSV40 | BPSV40-NLP-3xHA-SV40 | 0.53 ±0.01 |
| BPSV40 | BPSV40-SV40 | 0.73 ±0.07 |
| BPSV40 | BPSV40-SV40-EFE-NLP | 0.81 ±0.09 |
| BPSV40 | BPSV40-vBPSV40-EFE-NLP | 0.74 ±0.01 |
| BPSV40 | BPSV40-vBPSV40 | 0.69 ±0.05 |
| BPSV40 | BPSV40-vBPSV40-EFE-BPSV40 | 0.70 ±0.01 |
| BPSV40 | BPSV40-NLP-3xHA-SV40 | 0.37 ±0.05 |
| vBPSV40 | BPSV40-SV40 | 1.02 ±0.11 |
| vBPSV40 | BPSV40-SV40-EFE-NLP | 0.64 ±0.04 |
| vBPSV40 | BPSV40-vBPSV40-EFE-NLP | 0.85 ±0.10 |
| vBPSV40 | BPSV40-vBPSV40 | 0.66 ±0.08 |
| vBPSV40 | BPSV40-vBPSV40-EFE-BPSV40 | 0.76 ±0.03 |
| vBPSV40 | BPSV40-NLP-3xHA-SV40 | 0.54 ±0.07 |

**Table 5:** Nuclear localization signals used in this study.

| Name | Type | Amino acid sequence | Rel. editing (mean ±S.D.) |
|---|---|---|---|
| SGGS-XTEN-SGGS | Flexible | SGGSSGSETPGTSESATPESSGGS | 1.0 ±0.09 |
| GGS x 4 | Flexible | GGSGGSGGSGGSS | 1.09 ±0.06 |
| GGS x 5 | Flexible | GGSGGSGGSGGSGGSS | 1.07 ±0.06 |
| GGS x 7 | Flexible | GGSGGSGGSGGSGGSGGSGGSS | 0.91 ±0.02 |
| GGS x 8 | Flexible | GGSGGSGGSGGSGGSGGSGGSGGSS | 0.92 ±0.02 |
| Long 1 | Flexible | SGGSSGGSSGSETPGTSESATPESSGGSSGGSSKSGGSSGGSS | 1.08 ±0.1 |
| Long 2 | Flexible | SGGSSGGSSGSETPGTSESATPESSGGSSGGSSKSGGSSGGSSK SGGSSGGSS | 1.04 ±0.11 |
| Long 3 | Flexible | SGGSSGGSSGSETPGTSESATPESSGGSSGGSSKSGGSSGGSSK SGGSSGGSSKSGGSSGGSS | 1.10 ±0.04 |
| Long 4 | Flexible | SGGSSGGSSGGSSGGSSKSGSETPGTSESATPESSGGSSGGSSK SGGSSGGSSKSGGSSGGSSKSGGSSGGSS | 1.07 ±0.04 |
| X-ten | Rigid | SGSETPGTSESATPES | 0.78 ±0.01 |
| Linker A | Rigid | GSAGSAAGSGEF | 1.13 ±0.04 |
| KL(A/E)AA | Rigid | KLAEAA | 1.11 ±0.06 |
| KL(A/E)AA x 2 | Rigid | KLAEAAAKEAAAKAAA | 1.04 ±0.01 |
| KL(A/E)AA x 3 | Rigid | KLAEAAAKEAAAKEAAAKAAA | 1.04 ±0.03 |
| KL(A/E)AA x 4 | Rigid | KLAEAAAKEAAAKEAAAKEAAAKAAA | 1.05 ±0.07 |
| KL(A/E)AA x 5 | Rigid | KLAEAAAKEAAAKEAAAKEAAAKEAAAKAAA | 1.03 ±0.06 |
| SpyTag | Multimer | GGGSGSGLQAHIVMVDAYKPTKGGSGSGLQAHIVMVDAYKPTK | 0.16 ±0.02 |
| SunTag | Multimer | LLPKNYHLENEVARLKKLVGERGGSGGLLPKNYHLENEVARLKK LVGERGGSGGLLPKNYHLENEVARLKKLVGERGGSGGLLPKNYH LENEVARLKKLVGERGGSGGLLPKNYHLENEVARLKKLVGERGG SGGLLPKNYHLENEVARLKKLVGERGGSGGLLPKNYHLENEVAR LKKLVGERGGSGGLLPKNYHLENEVARLKKLVGERGGSGGLLPK NYHLENEVARLKKLVGERGSGEGRGSLLTCGDVEENPGPEDPPV ATMVMGPDIVMTQSPSSLSASVGDRVTITCRSSTGAVTTSNYAS WVQEKPGKLFKGLIGGTNNRAPGVPSRFSGSLIGDKATLTISSL QPEDFATYFCALWYSNHWVFGQGTKVELKRGGGGSGGGGSGGGG SSGGGSEVKLLESGGGLVQPGGSLKLSCAVSGFSLTDYGVNWVR QAPGRGLEWIGVIWGDGITDYNSALKDRFIISKDNGKNTVYLQM SKVRSDDTALYYCVTGLFDYWGQGTLVTVSS | 0.10 ±0.01 |
| T2A | split | EGRGSLLTCGDVEENPGPEDPPVATMV | 0.85 ±0.04 |

**Table 6:** Linker sequences used in this study.

### Retron msr-msd sequences used in this study

Legend: homology arms; insert; msr; msd ; native msd

#### Efe1-RT

##### Native msr-msd sequence

Efe1:

```
  1 CGCCAGCAGT GGCAATAGCG TTTCCGGCCT TTTGTGCCGG GAGGGTCGGC GAGTCGCCGA CTTAACGCCA
 71 GTAGTTTGTC TATATACCCA AAGCCGCTTC ATTGTACTTA AGTACGCTTT GCGTACGTCG CGCTGACGCG
141 CTCAGTACAG TTACGCGCCT TCGGGATGGT TTGATGGTAT TGCCGCTGTT GGCG
```

Eco8:

```
  1 GCCAGCAGTG GCAATAGCGT TTCCGGCCTT TGTGCCGGGA GGGTCGGCGA GTCGCCGACT TAACGCCAGT
 71 AGTATGTCCA TATACCCAAA GTCGCTTCAT TGTACCTGAG TACGCTTCGC GTACGTCGCG CTGACGCGCT
141 CAGTACAGTT ACGCGCCTTC GGGATGGTTT AATGGTATTG CCGCTGTTGG C
```

Vap1:

```
  1 CGCACCCTTA GCGAATGAGC TTACTTAGTT CATTGGATAG CGTTTCGCTA TCCTGCATAC AATCTGATTC
 71 AATGCCGCAT GAAAATGTGC AGAGCCAGAA TACAGTAGTT TCTGGAACTG CACATTTTCA TCCGCGACTT
141 AAGACGTAAG GGTGTG
```

Cex1:

```
  1 AGTGGTCGAG AGAGGTCTGG ACCGCATCAG CCTTAACGCC TCGAGCGTAG GAACGGCGTT GCGCCGTTCT
 71 GGTTGAAATG CTGGACACTC TCCGCAAGGT AGCCTGTTCT TGGCTCTCTC CCTCCCGAGC ACTACCGTCG
141 GGGTGGGAAG CGGAACCAAC GACGCAGCCG CCGTTTTCCC ACCCCGACGG TAGTGCTCGG GAGGGGAGAG
211 CCGGTGAGGC TACCGTGCCC CAGGTGAGCT GGTGGTGCCT TCCTGGCCTC CCTCGACCGC TCGC
```

Vro1:

```
  1 CACACCCTTA GCGAATGAGC TAACTTAGTT CATTGGATAG CGTTTCGCTA TCCTGCATAC AATCTGATTC
 71 AATGCCGCAT GAAAATGTGC AGAGCCAGAA TACAGTCGTT TCTGGAACTG CACCTTTTCA TCCGCGACTT
141 AAGACGTAAG GGTGTG
```

##### RFP reporter

```
  1 TCGGGGATGC CCTGGGTGTG GTTGATGAAG GCTTTGCTGC CGTACATGAA GCTGGTAGCC AGGATGTCGA
 71 AGGCGAAGGG G
```

##### Genomic loci in HEK293T cells

EMX1:

```
  1 TGGCCTGCTT CGTGGCAATG CGCCACCGGT TGATGTGATG GGAGCCCTTC GCTACATGCT TTCTTCTGCT
 71 CGGACTCAGG CCCTTCCTCC TCCAGCTTCT GCCGTTTGTA
```

AAVS1:

1 GGAGACTAGG AAGGAGGAGG CCTAAGGATG GGGCTTTTCT GTCACCAATC GCTACATGCT CTGTCCCTAG
71 TGGCCCCACT GTGGGGTGGA GGGGACAGAT AAAAGTACCC

HBB:

1 CTGCCCAGGG CCTCACCACC AACTTCATCC ACGTTCACCT TGCCCCACAG GCTACATGCT GGCAGTAACG
71 GCAGACTTCT CCTCAGGAGT CAGATGCACC ATGGTGTCTG

CFTR:

1 TATAAAAAGA TTCCATAGAA CATAAATCTC CAGAAAAAAC ATCGCCGAAG GCTACATGCT GGCATTAATG
71 AGTTTAGGAT TTTTCTTTGA AGCCAGCTCT CTATCCCATT

F9:

1 GTGAACATGA TCATGGCAGA ATCACCAGGC CTCATCACCA TCTGCCTTTT GCTACATGCT AGGATATCTA
71 CTCAGTGCTG AATGTACAGG TTTGTTTCCT TTTTTAAAAT

BRD8:

1 GGAGACTAGG AAGGAGGAGG CCTAAGGATG GGGCTTTTCT GTCACCAATC GCTACATGCT CTGTCCCTAG
71 TGGCCCCACT GTGGGGTGGA GGGGACAGAT AAAAGTACCC

**Genomic loci in U2OS**

TOMM70A:

1 AGTGTCTTTA GGGTTCAGTT GAAGAGGGGG TAAACTTTTA AAAAGAGGGT CAGTCTGCTT TCCCCCTGTT
71 TTATGTAATC CCAGCAGCAT TTACATACTC ATGAAGGACC ATGTGGTCAC GGCCGCCACC TAATGTTGGT
141 GGTTTTAATC CGTATTTCTT TGCAACTTCT GTCTGGGCAT GGGCGGCATC GCAAAGTGAA

RAB11:

1 TAGAGTGCGA GAGCCCATGG CCTCACCTTT AAAGAGGTAG TCGTACTCGT CGTCCCGTGT GCCGCCGCCA
71 CCTGTAATCC CAGCAGCATT TACATACTCA TGAAGGACCA TGTGGTCACG CATTGCGCGG CCGAGGAGCG
141 AAAGGGCGGG AGCAGCAGTG GTATCTGTGG GACCAGGGGG CGTCGCTGCA GGGGTAACCC

**Correcting Kif6 mutations**

1 TTACCGCCCA AACTGTCTCT GAGTACAGAG GTCATCATTG AATTCCTATA GGGGATGTGG GACCTATCCT
71 TCTCTGACAG GGCTATGATG ACCTAAACCA
